# Dissecting RNA Selectivity Mediated by Tandem RNA-Binding Domains

**DOI:** 10.1101/2024.10.17.618930

**Authors:** Sarah E. Harris, Yue Hu, Kaitlin Bridges, Bryan B. Guzmán, Francisco F. Cavazos, Justin G. Martyr, Jernej Murn, Maria M. Aleman, Daniel Dominguez

## Abstract

RNA-protein interactions are pivotal to proper gene regulation. Many RNA-binding proteins possess multiple RNA-binding domains; however, how these domains interplay to specify and regulate RNA targets remains poorly understood. Here, we investigate three multi-domain proteins, Musashi-1, Musashi-2, and Unkempt, three factors which share a high degree of RNA specificity. We use a combination of massively parallel *in vitro* assays with random or naturally derived RNA sequences and find that individual domains within a protein can have differing affinities, specificities, and spacing preferences. Further, we emphasize that while all three proteins have overlapping motif specificities, non-overlapping sequences may allow for target discrimination. We carry out large scale competition assays between these proteins and determine how individual protein specificities and affinities influence competitive binding. Integration of *in vivo* binding and regulation with *in vitro* specificities shows that target selection involves a combination of the protein intrinsic specificities described here, but cellular context is critical to drive these proteins to motifs in specific transcript regions. Finally, evolutionarily conserved RNA regions display evidence of binding multiple RBPs *in vivo*, and these RNA regions recapitulate this trend with the highest affinity *in vitro*. We highlight the importance of understanding features of complex RNA-protein interactions and how protein-target discrimination can be established.

## Introduction

Post-transcriptional gene regulation is a crucial step of gene expression. This regulation is primarily directed by RNA-binding proteins (RBPs) which bind and control RNA processing from synthesis, through maturation and degradation^1–4^. Over 1,500 RBPs have been discovered to date within the human proteome, many of which are comprised of multiple RNA-binding domains (RBDs) like RNA recognition motifs (RRMs), K homology (KH) domains, and zinc fingers (ZnFs)^5^. These RBDs serve to define RNA sequence preferences and often work in tandem to enhance avidity and affinity to a given RNA^6–8^. Whereas single domain RBPs are thought to bind weakly to short sequence motifs, two or more RBDs in tandem can greatly enhance the strength and surface area of the interaction, increasing sequence specificity and affinity^6, 7, 9–16^.

Studies on the evolution and conservation of RBPs and RBDs (reviewed by Gerstberger et al.^17^) reveal genomic interplay between domains. A large co-occurrence of highly structured RBDs with more degenerate low complexity domains (LCDs)^5^ within an RBP gene may allow for structured domain duplication at and around the LCD as the DNA sequence encoding LCDs is less stable than for structured domains, lending to local DNA expansions during replication^18,19^. Further, the evolutionary timeline in which tandem domains evolved provides further insights into their functional importance. For RRMs, duplications are frequently observed in the mammalian lineage and are more similar within RBPs than across, whereas KH domains evolved much earlier (with many KH domain proteins conserved through *S. cerevisiae*) with higher levels of RBD conservation across genes rather than within^9,17,20,21^. These differences may relate to RNA regulatory expansions where mammals have evolved complex alternative splicing reliant on a diverse set of RBPs^17,22,23^. Still largely unknown, however, are how these multi-domain RBPs interact with RNA from a quantitative biophysical standpoint: How do tandem domains lead to sequence targeting and enhanced affinity? Are the affinities of individual RBDs within a protein similar? Does the ordering of RBDs within a protein impact binding?

At a trans (protein-guided) level, many *in vitro* and *in vivo* studies have sought to define the RNA-binding capabilities for a large portion of the RBP family (selected large-scale studies: Pantier et al.^24^, Jolma et al.^25^, Van Nostrand et al.^26^, Dominguez et al.^27^, Van Nostrand et al.^28^, Ray et al.^29^). These studies largely found that RBPs tend to bind short (3-8 nucleotides) and relatively unstructured motifs within a given RNA. However, when examining the sequence preferences across RBPs, little motif diversity exists, where many RBPs exhibit similar RNA motif preferences^27^. A high degree of amino acid conservation between RBDs can be used as a proxy for predicting specificity where RBDs with >70% conservation are likely to have highly similar RNA motif preferencesc^29^. However, even when only proteins with differing RBD types are considered, RBPs as a group still appear to have a high degree of overlapping specificity^27^.

Examining the cis (RNA sequence) side, short motifs are insufficient to define *in vivo* binding as they can be found in nearly every transcript (reviewed by Achsel and Bagni^8^): And if every RBP bound to every occurrence of its motif within the transcriptome, regulatory diversity would be challenging. Instead, when a motif is present within the transcriptome, it is bound by a given RBP on average less than 10% of the time^8,27,30^. For example, the UGCAUG motif previously defined for the highly conserved alternative splicing factor RBFOX2^31^ can be found in close to 90% of transcripts^8^ but only appears to be bound by RBFOX2 in less than 7%^31^. Similarly, for the important neuronal protein Unkempt (UNK), the defined consensus motif UAGNNUUU is bound just over 20% of the time within the transcriptome^11^. Further, several RBPs have been shown to bind to regions not defined by a canonical motif. For example, UNK’s consensus motif only accounts for 15% of the protein’s total binding sites highlighting flexibility in target selection^11^. Therefore, from an RNA-centric view, RNA-protein interactions are not solely governed by primary RNA motifs: Rather surrounding sequence context and RNA secondary structure also play a pivotal role^26,27,29,32,33^.

In this manner, RNA binding is far from straightforward. Not only are individual RNA-protein interactions critical within every step of RNA processing, but other cellular factors influence aspects of these interactions. While previous studies have shown that *in vitro*-derived motifs correlate with *in vivo* binding data^29,30,33^, additional factors are known to contribute to target selection. Protein-protein or protein-ion interactions serve to either recruit or hinder specific RNA-protein interactions, which can lead to different *in vivo* binding patterns than would be expected based on *in vitro* affinities^32^. This culminates to RNA-protein interactions being impacted by i) expression levels of the RNA and protein of interest, ii) competitor proteins, iii) local ion concentrations, iv) diffusion, and a host of other factors.

To begin to fully understand how RBPs interact with RNA, we must first understand how tandem RBDs work together—or in competition—to define an RNA target. Both full-length RBPs as well as individual RBDs have been shown to interact with 3 to 8 nucleotide sequences^26–28,34^. This view oversimplifies the reality of RNA-protein interactions and must be expanded. How do RBPs select for a given sequence, and can their diversity of *in vivo* interactions be attributed to differences in RBDs for multi-domain RBPs? In the present study, we sought to explore the RNA-binding preferences of three RBPs: Musashi-1 (MSI1), Musashi-2 (MSI2), and UNK. These proteins have all been either directly shown or indirectly implicated to bind a UAG core motif^11,27,35–40^. However, each have distinct functions where MSI1 and MSI2 are important in development and have been shown to drive cancer progression^41–49^ (reviewed by Forouzahfar et al.^50^) and UNK mediates neuron morphology^11^. Further, all three proteins are involved in translational regulation^51–53^. MSI1 and MSI2 are comprised of two RRMs whereas UNK has six ZnFs which serve as the primary mediators of RNA interactions^35,36,39,54^. Here, we examine these proteins domain-by-domain to determine the nature of each domain’s RNA-binding capabilities as well as define how each domain contributes to overall specificity and selectivity for the given protein. We explore the competition potential of two of these proteins—MSI1 and UNK—and find that RBP-specific binding patterns rely on a combination of primary motif strength and multi-valent interactions. These studies highlight the cross-regulatory potential of RBPs while also defining their independent functionalities as dictated by individual RBDs.

## Results

### UAG-Binding Proteins have Similar Specificities

To understand how individual RBDs contribute to affinity, we nominated three multi-domain RBPs with similar RNA sequence specificity: MSI1, MSI2, and UNK (**Figure 1A**). To test the similarity of binding preferences between these tandem domain-containing proteins, we utilized RNA bind-n-seq (RBNS), an unbiased massively parallel *in vitro* assay^27,55^. The strength of RBNS is that it allows for a quantification of the effects of RNA sequence, secondary structure, and surrounding sequence context in a single assay, by allowing proteins to select binding partners from millions of RNAs in parallel^27,55^. Additionally, RBNS does not rely on chemical or light-based crosslinking of protein to RNA such as CLIP-based methods, and therefore is not impacted by crosslinking biases (reviewed by Chakrabarti et al.^56^). RBNS, instead, relies solely on biochemical RNA-protein interactions where binding reactions are carried out between recombinant protein and a diverse RNA pool, RNA-protein complexes are isolated, bound RNA is sequenced, and sequence enrichments of protein-associated RNA are determined in relation to an input RNA pool^55^.

**Figure 1.**
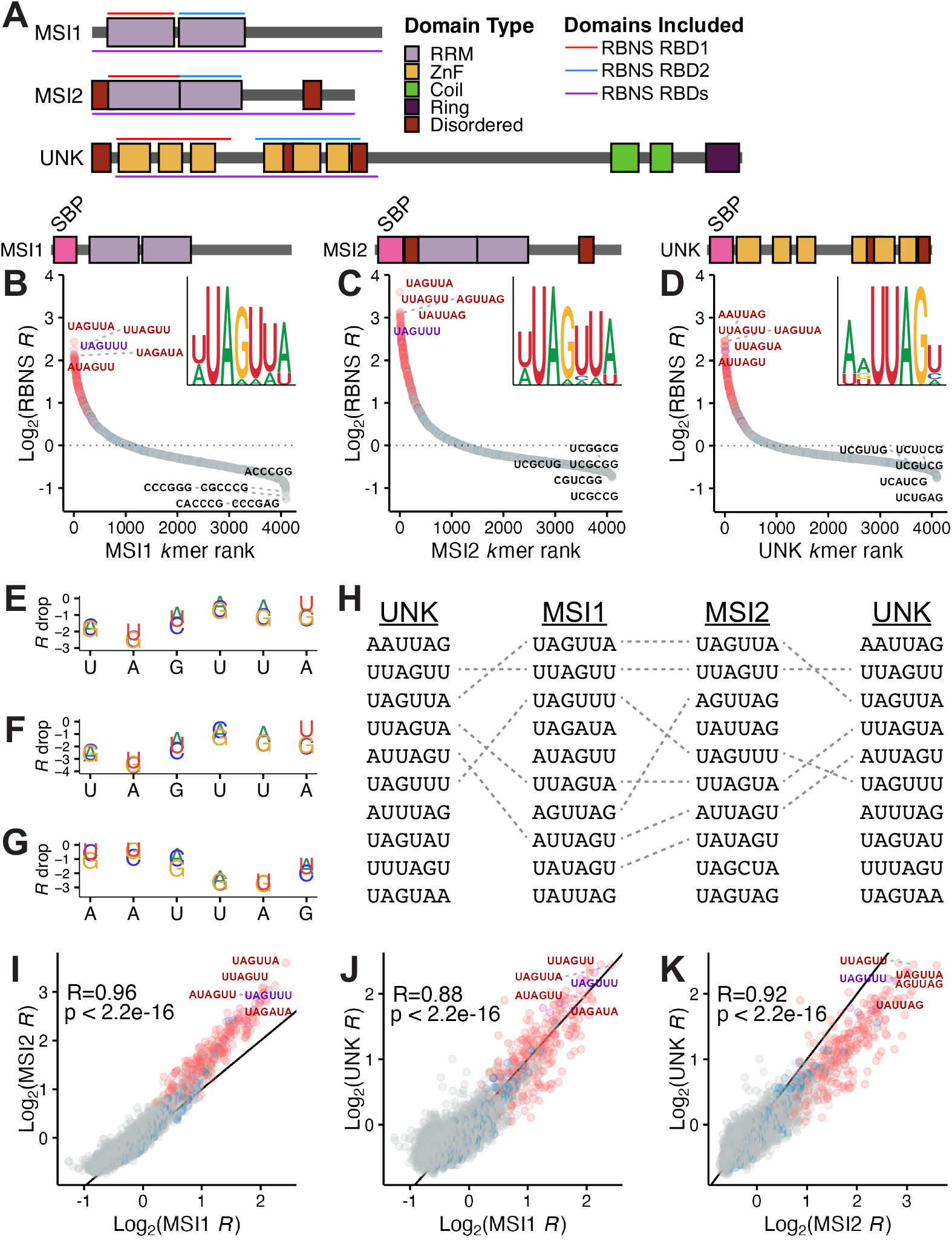
Motif preferences for full-length MSI1, MSI2, and UNK as determined via RBNS. A) Domain architecture of MSI1, MSI2, and UNK as comprised of RRMs (light purple), ZnFs (yellow), coiled domains (green), ring domains (dark purple), and disordered regions (dark red). Regions used in individual domain RBNS experiments are highlighted by red and blue lines where regions used in full-length RBNS experiments are highlighted by purple lines. B-D) Scatter plot of all 6mer log_2_ enrichments for B) MSI1, C) MSI2, and D) UNK as determined via RBNS. MSI1 and UNK assayed by Dominguez et al.^27^. 6mers colored by presence of “UAG” (red), “UUU” (blue), both “UAG” and “UUU” (purple), or none (grey). Logo was generated from the top 15 6mers. E-G) Relative enrichment plot of each nucleotide within the top 6mer for E) MSI1, F) MSI2, and G) UNK. R drop was calculated as the change in enrichment from the top 6mer upon mutation of a single nucleotide. H) Overlap of the top ten 6mers for MSI1, MSI2, and UNK. I-K) Correlation plot of I) MSI1 versus MSI2, J) MSI1 versus UNK, and K) MSI2 versus UNK log_2_ enrichments. MSI1 and UNK data obtained from GEO^27^. 6mers colored by presence of “UAG” (red), “UUU” (blue), both “UAG” and “UUU” (purple), or none (grey). Pearson’s correlation coefficient and p value included.

All three proteins have been demonstrated to bind to a UAG core motif *in vitro* or *in vivo*^11,27,35–40^. To confirm this, we obtained RBNS data from ENCODE for MSI1 and UNK^27^ and individually assayed MSI2. With these data, we confirmed the findings of *in vivo, in vitro*, and crystallography studies^11,27,35–40^ showing that all three proteins bind a UAG core motif with high specificity (**Figure 1B-D & Supp. Figure 1A**). Because RBNS data provides a full spectrum of enrichment values (a proxy for affinity^55^) for all possible *k*mers, we calculated the relative importance of each position for the top 6mer by changing a base at each position and determining the consequence on enrichment (*R* depletion).

For all three proteins, disruption of the UAG had the greatest impact on binding where upstream and downstream flanking sequences exhibited less severe penalties in binding enrichments (**Figure 1E-G**). The least severe change at the UAG was to UAA or UAU consistent with data presented below.

To determine the degree of shared specificity, we overlapped the top ten 6mers across these proteins and observed at least 50% are shared between proteins (**Figure 1H**). Of the top 25 bound 6mers across the three proteins, on average 20 overlapped between two and 15 overlapped between all three. The degree of overlapping specificity was also reflected by Pearson’s correlation coefficients with all three sets of RBPs being highly correlated in *k*mer enrichments (MSI1 vs. MSI2: R=0.96, MSI1 vs. UNK; R=0.88, and MSI2 vs. UNK: R=0.92) (**Figure 1I-K**). These data confirm that MSI1, MSI2, and UNK all prefer a UAG core motif, tolerate similar changes to that motif, and emphasize their potential for binding site overlap.

### Individual Domains of UAG-Binding Proteins Contribute to Protein Individuality

Despite the top motifs for MSI1, MSI2, and UNK having a UAG-core, we hypothesized that the individual domains may have differing preferences across proteins and therefore allow for protein-specific target selection. To test this, we separated each protein into its individual RNA-binding units—two RRMs for MSI1 and MSI2 and two ZnF clusters (ZnF1-3 and ZnF4-6) for UNK^54,57^—and performed RBNS^27,55^with these individual domains. We also utilized datasets we previously generated for UNK ZnF clusters^33^, and replicated those RBNS assays for this study.

From this, we found that both individual RRMs for MSI1 and MSI2 preferentially bind a UAG-core motif (**Figure 2A-B, D-E & Supp. Figure 2A-B**), consistent with previous reports and our own data^27,36–39^(**Figure 1B-C**). Of the top 25 6mers, 16 were cross-bound between MSI1’s RRMs and 14 were cross-bound for MSI2’s RRMs. As a metric of conservation versus RNA-binding patterns, we compared the individual RNA-binding capabilities of the RRMs of MSI1 and MSI2 paralogs. When looking at intra-protein binding enrichments, we observe that MSI1’s RRMs are more highly correlated than those of MSI2 (R=0.93 and R=0.86, respectively) (**Figure 2C, F**). Previous work has demonstrated that greater than 70% amino acid conservation can be predictive of binding specificity^29^. However, while the binding preferences for MSIs’ RRMs are highly correlated, RRM1 and 2 conservation is only 45% for MSI1 and 42% for MSI2. To understand this relatively low conservation but high binding similarity we looked at an available NMR structure for MSI1 RRM2 in complex with RNA^36^, and found that all RNA-contacting residues^36^ were conserved between domains^58^, save two which shared chemical similarity (as predicted by BLAST, D91 in RRM1 to E180 in RRM2 and R99 in RRM1 to K183 in RRM2). Further, between several RNA-contacting residues, a 5 amino acid stretch exists in RRM1 that is not found in RRM2 (**Supp. Figure 2D**). These differences in combination may define the individual RNA-binding profiles for the two domains or may impart affinity differences (discussed in more detail below). For example, RRM1 prefers 6mers where the UAG is flanked upstream by A/U whereas RRM2 prefers G/U flanking nucleotides. Using amino acid sequence conservation to predict specificity remains a challenging task and conservation of protein structure and RNA contacts can overcome overall low amino acid conservation.

**Figure 2.**
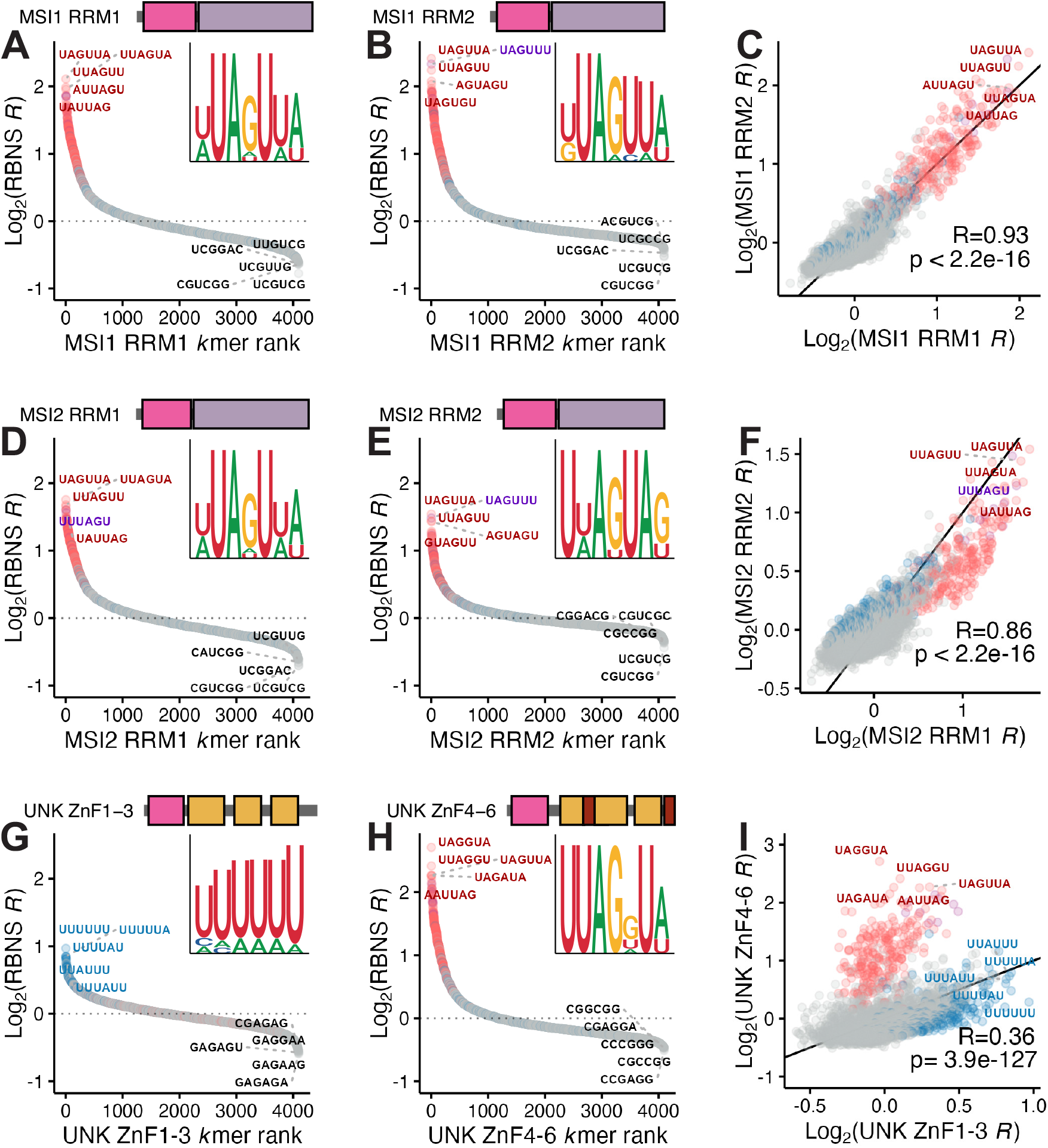
Motif preferences for individual RBDs of MSI1, MSI2, and UNK as determined via RBNS. A-B) Scatter plot of all 6mer log_2_ enrichments for MSI1 A) RRM1 and B) RRM2 as determined via RBNS. 6mers colored by presence of “UAG” (red), “UUU” (blue), both “UAG” and “UUU” (purple), or none (grey). Logo was generated from the top 15 6mers. C) Correlation plot of MSI1 RRM1 versus MSI1 RRM2 log_2_ enrichments. 6mers colored by presence of “UAG” (red), “UUU” (blue), both “UAG” and “UUU” (purple), or none (grey). Pearson’s correlation coefficient and p value included. D-E) Scatter plot of all 6mer log_2_ enrichments for MSI2 D) RRM1 and E) RRM2 as determined via RBNS. 6mers colored by presence of “UAG” (red), “UUU” (blue), both “UAG” and “UUU” (purple), or none (grey). Logo was generated from the top 15 6mers. F) Correlation plot of MSI2 RRM1 versus MSI2 RRM2 log_2_ enrichments. 6mers colored by presence of “UAG” (red), “UUU” (blue), both “UAG” and “UUU” (purple), or none (grey). Pearson’s correlation coefficient and p value included.G-H) Scatter plot of all 6mer log_2_ enrichments for UNK G) ZnF1-3 and H) ZnF4-6 as determined via RBNS. Replicate data from this study was merged with that previously reported by Harris et al.^33^. 6mers colored by presence of “UAG” (red), “UUU” (blue), both “UAG” and “UUU” (purple), or none (grey). Logo was generated from the top 15 6mers. I) Correlation plot of UNK ZnF1-3 versus UNK ZnF4-6 log_2_ enrichments. Replicate data from this study was merged with that previously reported by Harris et al.(Harris et al. 2024). 6mers colored by presence of “UAG” (red), “UUU” (blue), both “UAG” and “UUU” (purple), or none (grey). Pearson’s correlation coefficient and p value included.

For UNK, we observed that ZnF1-3 preferentially binds an A/U rich motif whereas ZnF4-6 associates with the primary UAG-core (**Figure 2G-H & Supp. Figure 2C**). We and others have previously observed these patterns with these domains^11,27,33,35^. Due to these separate preferences between the individual domains, their binding patterns are not highly correlated where A/U rich motifs are top bound for ZnF1-3 but poorly enriched for ZnF4-6 with the opposite being true for UAG-core motifs (**Figure 2I**). Of the top 25 bound 6mers for each domain, none of them are cross-bound between ZnF1-3 and ZnF4-6. UNK’s ZnF clusters are 32% conserved, yet their binding preferences differ drastically. Interestingly, when aligning these domains with BLAST^58^, little sequence conservation exists across domains, however, the key RNA-contacting residues as defined with crystal structures^35^ are positionally conserved, but largely not amino acid conserved (**Supp. Figure 2E**). These data show how individual RBDs have differing motif preferences than the full-length RBP.

### Spacing and Contextual Preferences Varies for UAG-Binding Proteins

How tandem RBDs work together to bind longer RNA targets can be complex and difficult to define experimentally. While we have highlighted the top motif for each RBD and full-length protein, as can be seen above, there are many short 3mers that are well-bound by each RBD (**Supp. Figure 2A-C**). The affinity of each RBD to these sequences, the spacing of the motifs, and the structural context in which they are embedded are all expected to generate a large repertoire of targets with similar affinities. Previous studies investigating the paralogs heterogeneous nuclear ribonucleoprotein L (hnRNPL) and hnRNPL-like (hnRNPLL) found that both RBPs bind the same *k*mers in a bipartite manner (e.g. split motif), but their spacing preferences are different^59^. Therefore, we sought to investigate the spacing preferences for MSI1, MSI2, and UNK in an unbiased manner.

To address this question, we developed a version of RBNS (positional RBNS or posRBNS) where the preferred motif is locked centrally, and the surrounding nucleotides are randomized during DNA synthesis (Methods), resulting in very diverse pools of RNAs each with a central UAG. This approach increases our ability to determine complex motifs, motif spacing, and RNA secondary structure by enabling the analysis of millions of bound sequences all containing the UAG of interest. Using positional RBNS, we quantified the impacts of every motif upstream and downstream of the locked UAG and any preferred spacing between for each of these proteins (**Figure 3A**). For MSI1 and MSI2, as expected given our results above, the top secondary motif outside of the core-UAG motif is an additional UAG motif (**Figure 3B-C & Supp. Figure 3A, C**). To determine how other motifs might come into play we filtered our data such that only the central UAG was present (i.e. no other UAG anywhere else in the read were included) and asked what secondary motifs were enriched. UAA, UUA, and AUA were enriched downstream while GUA, UAA, and CCC were enriched upstream for the MSI paralogs. For UNK, we observed a downstream A/U rich preference with variable spacing between one and four nucleotides (**Figure 3D & Supp. Figure 3E**). Upstream of the locked UAG core, we observed strong preference for a C-rich motif, again with variable spacing (**Figure 3D & Supp. Figure 3E**). We have little evidence that these are directly bound by these RBDs and instead believe them to be modifiers of RNA secondary structure as discussed further below. While these data highlight the uniqueness of these proteins it also demonstrates their specificity overlap at both the primary and secondary motif levels.

**Figure 3.**
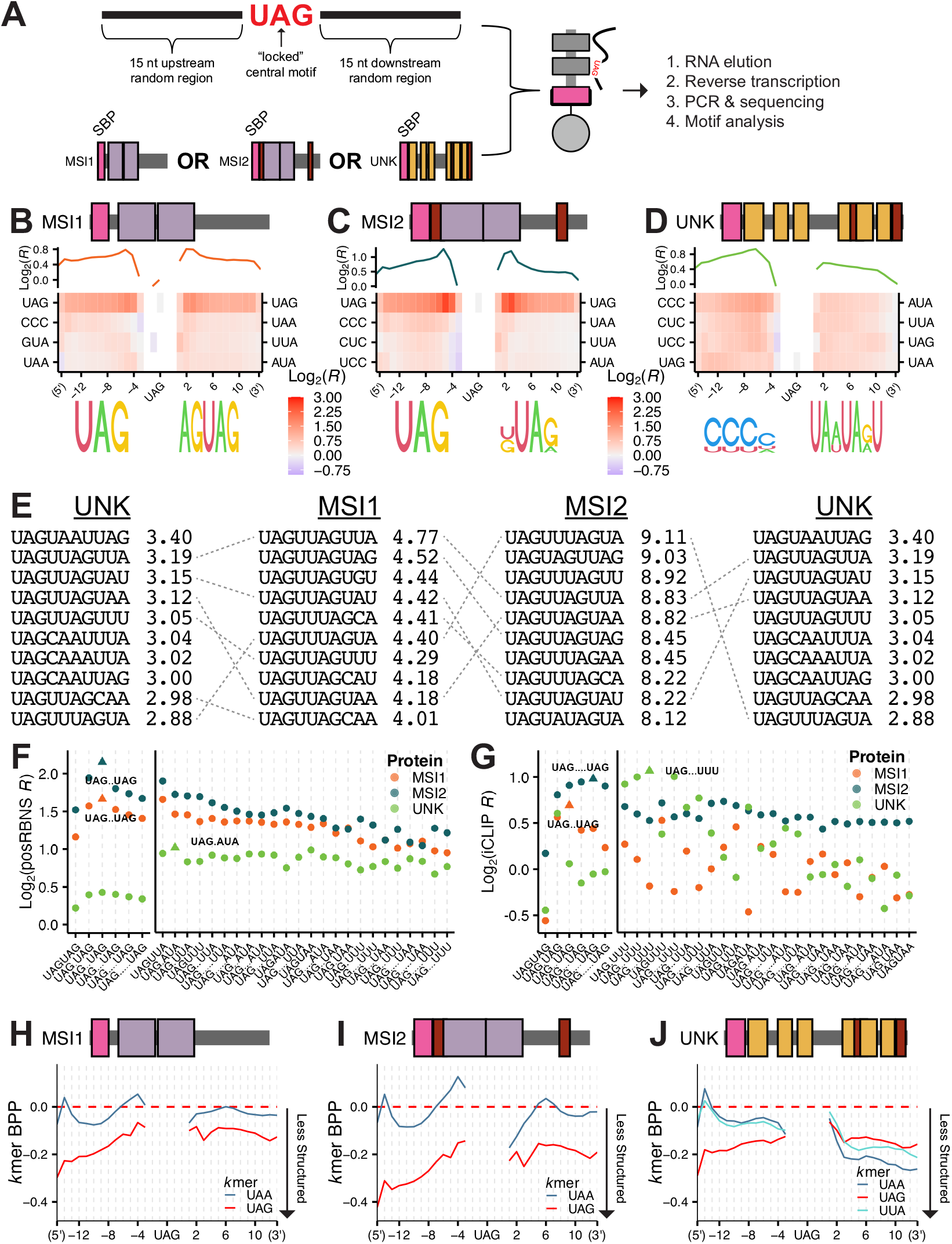
Secondary motif, spacing, and base pair probability preferences for MSI1, MSI2, and UNK as determined via posRBNS. A) Design of posRNA and layout of posRBNS. B-D) Heat map of the log_2_ posRBNS enrichment of top four enriched 3mers for B) MSI1, C) MSI2, and D) UNK. Logos were generated from the top five 4mers. Log_2_(R) line plot shows average enrichment of top four 3mers positionally. E) Overlap and enrichment of the top ten UAGN7 10mers for MSI1, MSI2, and UNK. posRBNS enrichment values shown. F) Log_2_ posRBNS enrichment of bipartite motifs for MSI1 (orange), MSI2 (blue), and UNK (green). Top bipartite motif and spacing denoted by triangle. G) Log¬2 iCLIP enrichment of bipartite motifs for MSI1 (orange), MSI2 (blue), and UNK (green). Top bipartite motif and spacing denoted by triangle. H-J) Line plot of the base pair probability of the top two enriched 3mers for H) MSI1, I) MSI2, and J) UNK.

Given the depth of this experiment, we were able to analyze 10mers starting with the locked UAG (UAGN_7_). In split motifs, intervening nucleotides are often denoted as Ns, but it is likely that preferences exist in these positions that strengthen interactions. This type of analysis has previously been a challenge given that there are 4^10^ ( 1e6) possible 10mers and quantifying the full spectrum of 10mer enrichments with robustness is a significant challenge as most experiments lack the depth. However, since we are most interested in UAG-containing 10mers and every sequence has a UAG, posRBNS is well suited for this analysis. Across all three proteins, 10mer analysis demonstrated the highest enrichments for UAGNUAG and UAGNNUAG where N was preferentially U/A, see figure for specific sequences (**Figure 3E**). Further, all three proteins exhibit strong binding patterns *in vitro* to UAGN_1-5_UAG and UAGN_1-5_(A/U-rich) (**Figure 3F**). Strikingly, even though there are a total of 16,000 UAG-containing 10mers in this experiment, the top 10mers were still largely shared for each RBP.

As noted by previous studies, lowly enriched *in vitro* motifs may be tighter bound *in vivo* due to protein-protein interactions at and surrounding the motif or increased RBP expression^32,60^. To test how these *in vitro* patterns compared to *in vivo* patterns, we examined available iCLIP data for all three proteins^11,37,38^. We identified iCLIP peaks for each protein, determined their transcript region, and selected “control peaks” matched for transcript region. Similar to the RBNS approach, we next computed motif enrichments within protein bound peaks relative to control peaks (see Methods). We observed similar enrichment patterns between nsRBNS and iCLIP where MSI1 and MSI2 preferentially bound to bipartite UAG motifs (with varied spacing) and UNK preferentially bound to UAGN_3_UUU (**Figure 3G**). While these patterns are slightly different from what was observed *in vitro, in vitro* patterns for each protein are still highly frequent *in vivo*. Further, we cannot exclude that crosslinking biases that generally favor Us impact the detection of motifs in iCLIP. These data highlight the power of unbiased *in vitro* assays like RBNS where top enriched motifs are frequently observed *in vivo*^30^.

To understand if any RNA structural preferences impact binding patterns, we performed *in silico* RNA folding (RNAfold^61^) and determined the base pair probability (BPP) of every 3mer in our posRBNS experiment. When looking at the top secondary motifs (UAG and UAA for MSI1 and MSI2; UAA, UAG, and UUA for UNK), we observe lower BPP downstream of the locked central UAG motif (**Figure 3H-J & Supp. Figure 3B, D, F**), likely contributing to increased binding due to the unstructured RNA preferences for many RBPs^25,27^. As noted above (**Figure 3B-D**), we observed C-rich motifs as upstream enriched for all three proteins. We hypothesized that this abundance may be due to the intrinsic nature of C to not pair with downstream A/U rich regions, thus promoting motif accessibility. We noted that CCC motifs largely have the lowest BPP of all *k*mers upstream of the central UAG (**Supp. Figure 3 B, D, F-I**). Taken together, these data demonstrate that all three proteins preferentially bind a UAG motif, but binding is stabilized and enhanced by additional motifs, preferential spacing, and increased motif accessibility.

### MSI1, MSI2, and UNK Demonstrate Similar Binding Affinities and Multi-Domain Avidity

To test and compare affinities across proteins, we used fluorescence polarization (FP) against a UAG-containing oligo. We previously showed via FP that UNK binds this RNA sequence at 40 nM^33^. When performing FP with MSI1 or MSI2, we observe high affinity for this sequence where MSI1 binds at 5 nM and MSI2 at 2 nM (**Figure 4A-B**). Globally, this affinity difference between MSI1 and MSI2 was also observable via RBNS where MSI2 exhibited higher enrichments than MSI1 (**Supp. Figure 1A**). To test how each domain contributes to this binding affinity, we assessed their constituent domains individually. When doing this previously with UNK, we observed that ZnF4-6 bound a UAG-containing oligo approximately 5-fold better than ZnF1-3 ( 0.4 vs 2.4 μM, respectively)^33^. Interestingly, we observed similar patterns for MSI1 and MSI2 where one RRM bound with greater affinity (more than 5-fold better) than the other (**Figure 4A-B**). These findings hold despite each domain demonstrating strong enrichments with UAG-containing *k*mers (**Figure 2A-B, D-E**). Additionally, we tested binding of MSI1, MSI2, and UNK as well as their individual domains against random sequence RNA (N_16_) to determine if the RNA backbone contributed significantly to binding. While all three full-length proteins demonstrated some binding potential at high protein concentrations, these binding affinities were drastically lower compared to UAG-containing RNAs (**Supp. Figure 4A-C**). These data present a potential “tandem anchoring” mechanism for multi-domain RBPs where one domain mediates primary contacts (large anchor) and demonstrates higher affinity for a given RNA sequence and the second domain serves to enhance avidity to the RNA target (stabilizing anchor), reflecting previous studies that have highlighted similar cooperative potential between RBDs^9,12–16^(reviewed by Lunde et al.^6^ and Auweter et al.^62^).

**Figure 4.**
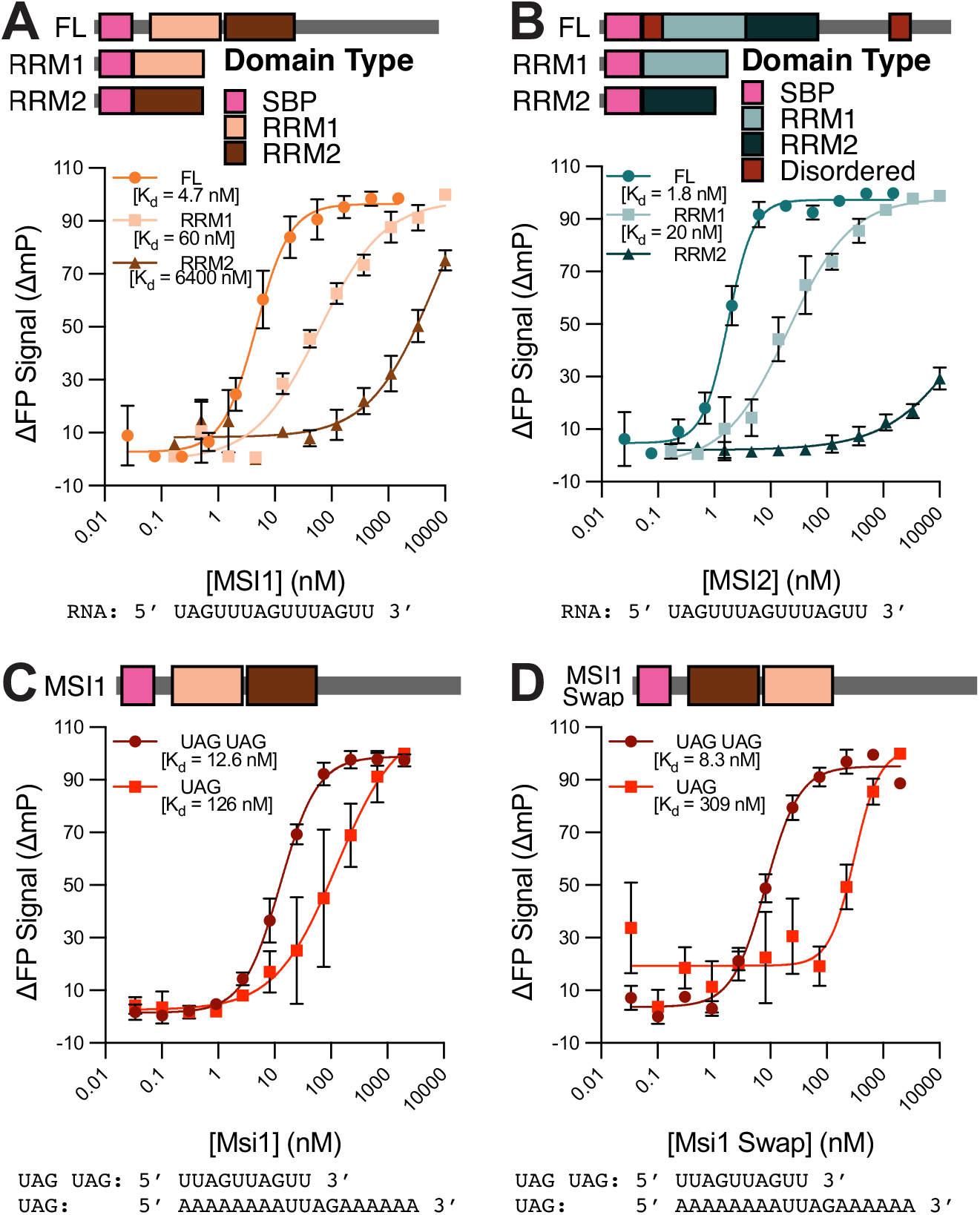
Affinity of MSI1 and MSI2 for UAG-containing sequences. A) Delta fluorescence polarization binding curves (n=3) for full-length MSI1 (orange circle), MSI1 RRM1 (light orange square), and MSI1 RRM2 (dark orange triangle) incubated with a tri-UAG-containing oligo. Full-length curve was normalized to its minimum and maximum fluorescence polarization signal with RRM1 and RRM2 normalized to the maximum of RRM1 to produce delta fluorescence polarization values. Data are presented as mean values +/-SD. B) Delta fluorescence polarization binding curves (n=3) for full-length MSI2 (teal circle), MSI2 RRM1 (light teal square), and MSI2 RRM2 (dark teal triangle) incubated with a tri-UAG-containing oligo. Full-length curve was normalized to its minimum and maximum fluorescence polarization signal with RRM1 and RRM2 normalized to the maximum of RRM1 to produce delta fluorescence polarization values. Data are presented as mean values +/-SD. C-D) Delta fluorescence polarization binding curves (n=3) for full-length C) MSI1 and D) MSI1 swap incubated with a mono-UAG-containing oligo (red square) or a dual-UAG-containing oligo (dark red circle). Each curve was normalized to its minimum and maximum fluorescence polarization signal to produce delta fluorescence polarization values. Data are presented as mean values +/-SD.

We further tested binding of MSI1 with varying combinations of UAG motifs. With a mono-UAG-containing oligo, we fit a K_d_ of 126 nM. However, the binding is enhanced 10-fold with the addition of just one UAG, likely accounted for by the dual-RBD architecture of MSI1 (**Figure 4C**). As both RRMs of MSI1 preferentially bind a UAG-core motif, we next asked whether orientation of these domains was important for MSI1 binding. We observe that by swapping the order of the domains, the affinity for a mono-UAG-containing oligo is decreased 2-fold. However, if the RNA is able to accommodate both domains (i.e. possesses a bipartite motif), no observable K_d_ difference is present upon domain swap (**Figure 4D**). We hypothesize that tandem RBD anchoring involving one strong and one weak RBD may be a mechanism that enables wider target selection and may be important for accommodating evolving motifs.

### Bipartite RNA Motifs Allow for Increased Affinity for Multi-Domain RNA-Binding Proteins

UNK is separate from MSI1 and MSI2 in that its “two” RBDs do not both prefer a UAG-containing sequence, but rather ZnF1-3 binds A/U rich RNA. Therefore, we designed fluorescent RNA oligos based on the findings from posRBNS (**Figure 3D**) to test the dual-domain avidity for UNK. UNK weakly binds to a mono-UAG oligo at 400 nM; however, adding in a secondary AUA enhanced binding affinity to 90 nM. As noted above, the presence of Cs was noted upstream of the UAG, surprisingly adding Cs upstream of the UAG enhanced affinity by 2-fold. However, Cs had little impact on binding affinity on oligos that contained both UAG and AUA. (**Figure 5A**). Despite these sequences being able to enhance binding, neither polyC nor polyU oligos demonstrated any binding on their own (**Supp. Figure 5A**). Thus, how Cs may act on these UNKs affinities remains unclear, but they are unlikely to make direct contacts with the ZnFs and even the most preferred secondary motifs (Us) on their own do not confirm high affinity binding.

**Figure 5.**
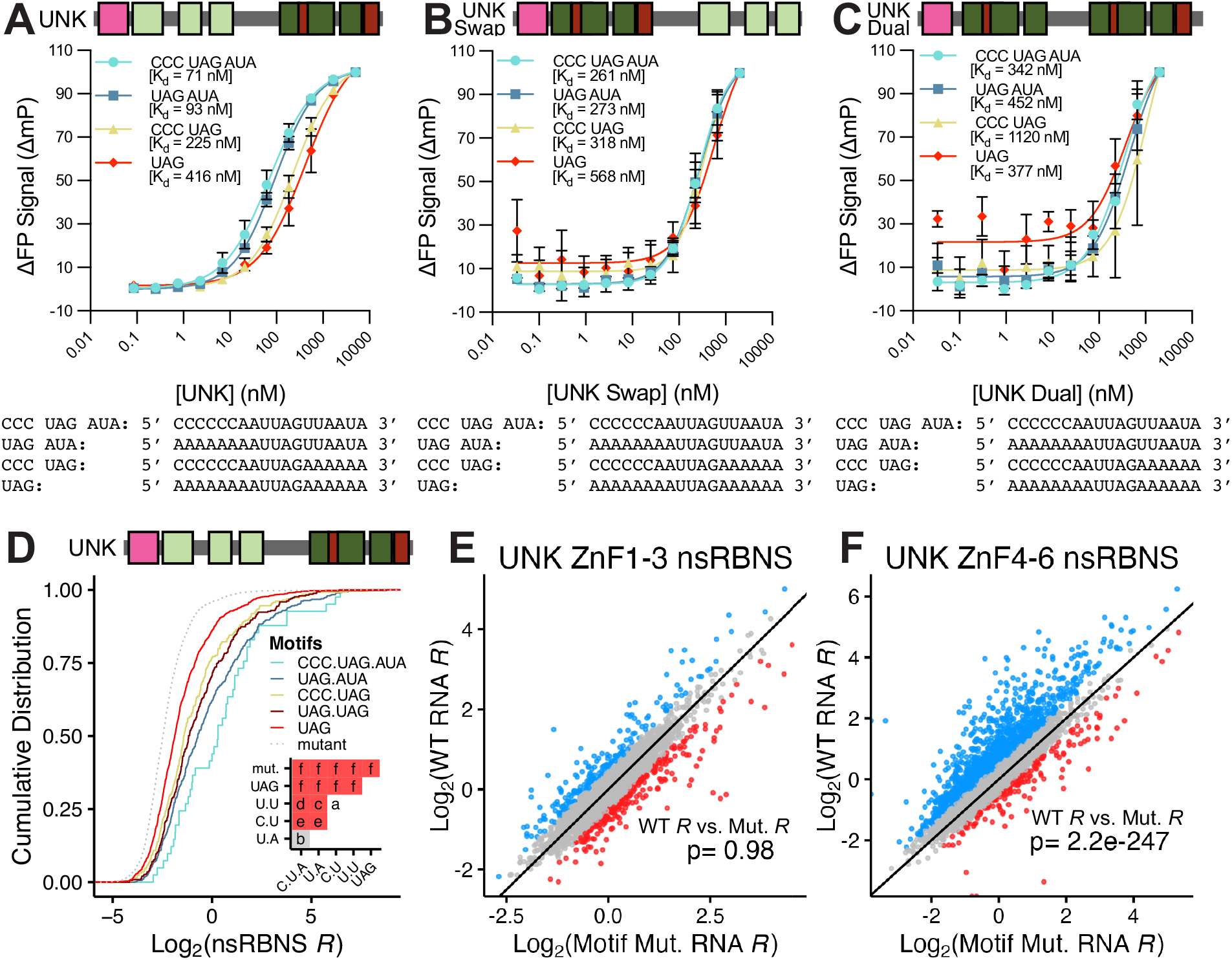
Combinatorial affinity of UNK for positional motifs. A-C) Delta fluorescence polarization binding curves (n=3) for A) UNK, B) UNK swap, and C) UNK dual incubated with a UAG-containing oligo (red diamond), CCC-UAG-containing oligo (yellow triangle), UAG-AUA-containing oligo (blue square), or a CCC-UAG-AUA-containing oligo (cyan circle). Each curve was normalized to its minimum and maximum fluorescence polarization signal to produce delta fluorescence polarization values. Data are presented as mean values +/-SD. D) Cumulative distribution function of log_2_ nsRBNS enrichments with UNK of positional-motif-containing oligos from Harris et al.^33^: CCC-UAG-AUA (cyan), UAG-AUA (blue), CCC-UAG (yellow), UAG-UAG (dark red), UAG (red), motif mutant (light grey, dotted). Inset shows significance values for all comparisons via two-sided KS test and corrected for multiple comparisons via the BH procedure. Red denotes significant (p<=0.05), and grey denotes nearing significant (p<=0.1). Values are as follows: a (ns), b (p<=0.1), c (p<=0.05), d (p<=0.01), e (p<=0.001), f (p<=0.0001). E-F) Scatter plot of log_2_ nsRBNS enrichments with E) UNK ZnF1-3 and F) ZnF4-6 of wildtype (Y-axis) versus motif mutant (X-axis) oligos. Log_2_ change in enrichment (wt-mut) was calculated for each sequence pair: > 0.5 defined as bound better in wt (blue), < -0.5 defined as bound better in mut (red), 0 +/-0.5 defined as similar binding (grey). Significance determined via paired, one-sided Wilcoxon test.

To test how domain order contributed to binding and further confirm each domain’s preferential sequences, we tested the same positional oligos against a swapped order domain construct of UNK (ZnF4-6, 1-3; UNK Swap). With this, we observed that a single UAG-containing RNA bound at 550 nM and adding secondary motifs CCC, AUA, or both enhanced this binding 2-fold (**Figure 5B**). However, this construct failed to recapitulate the full-strength binding of the canonically ordered domain and falls short by >3-fold. Further, we tested these sequences against a construct where ZnF1-3 were replaced with ZnF4-6 (UNK Dual, tandem ZnF4-6). When ZnF1-3 are not present, no secondary motif combinations had any impact on binding above a mono-UAG (**Figure 5C**). These data highlight the importance of domain order to RNA binding for tandem domain RBPs and show that UNK RBDs possess limited flexibility in that the preferred motif must contain a UAG followed by a secondary A/U rich motif.

We previously designed a natural sequence version of RBNS (nsRBNS) to test the effects of species-specific sequence differences on UNK binding patterns^33^. This pool was based on iCLIP data from human cells for UNK^11^. We identified bound regions in human and mouse and designed a natural sequence pool to unbiasedly test the binding of UNK to 25,000 natural (120 nt long) RNA sequences. We revisited this data to determine the enrichment of natural targets containing the above-described motifs and strong enrichment of UNK to CCC-UAG-AUA and UAG-AUA-containing sequences over a single UAG or two UAGs in tandem (**Figure 5D**). Confirming that these motifs embedded in longer natural sequence contexts enhance affinity, we included motif mutants where the central UAG was mutated to CCG. Considering these mutants where no UAGs were present, we see the largest depletion of the group.

To further highlight the differences between ZnF1-3 and ZnF4-6 to full length UNK binding against natural sequence, non-synthetic targets, we performed nsRBNS with the individual domains as previously described^33^. We rationalized that assaying against the individual domains would allow for an understanding of how each domain contributes global binding patterns observed for UNK *in vivo*. We hypothesized that ZnF1-3 would remain agnostic to motif mutants whereas ZnF4-6 would be largely impacted. Indeed, no large binding differences were observable for ZnF1-3 (**Figure 5E**) whereas ZnF4-6 largely displayed preferential binding to the wild-type oligos (**Figure 5F**).

To determine the effects of each cluster on overall binding patterns, we calculated the Pearson’s correlation coefficient for each domain versus full-length UNK in our nsRBNS assay^33^. We observe weak correlation for ZnF1-3 (R=0.27) and moderate correlation with ZnF4-6 (R=0.56) (**Supp. Figure 5B-C**). Further, when comparing ZnF1-3 with ZnF4-6, we observe weak correlation (R=0.44), continuing to highlight the differences between these domains (**Supp. Figure 5D**). These data demonstrate that while neither domain could recapitulate the full binding strength of the whole protein, ZnF4-6 is more similar, which is reflected in its preferred motif mirroring that of the full length (**Figures 1 & 2**), as well as its higher affinity for RNA^33^. Further, they show the sequence discrimination between ZnF1-3 and ZnF4-6 for UNK and highlight the combinatorial potential of multiple domains.

### MSI1 and UNK Compete for Some Binding Sites *in vitro*

Thus far, we have highlighted similarities and differences between MSI1, MSI2, and UNK and while some patterns are unique, we also show they bind identical motifs with high affinity (**Figures 1 & 4**). This raises the possibility that at times these proteins may compete for targets. Further, both MSI1 and UNK are expressed in the same cell types^63^. Therefore, we next wanted to determine how these proteins would behave in competition given their differing secondary motif preferences and affinities. To test this, we designed competition posRBNS (compPosRBNS) with the background of posRBNS as described above, but adding in competitor protein. This competitor protein (UNK for MSI1-centric or vice versa) was produced without an SBP tag, so only the assayed protein and bound RNAs would be captured with magnetic pulldown while the competitor titrates away RNA targets (**Figure 6A**). Starting with SBP-UNK, we titrated in 0, 0.1, 1, or 10X MSI1 and measured the effects of binding to UNK (**Figure 6B-D**). In this experiment, we observe that UNK-specific sequences—like upstream CCC (**Figure 6B**) or downstream AUA (**Figure 6C**)—are difficult to compete off with MSI1 and only see depletion at 10X competitor. We noted some apparent enhancement of CCC or AUA binding at lower MSI1 concentrations, which is likely expected in this assay^55^. Competition patterns still hold when only oligos with a single UAG motif are considered (**Supp. Figure 6A-B**).

**Figure 6.**
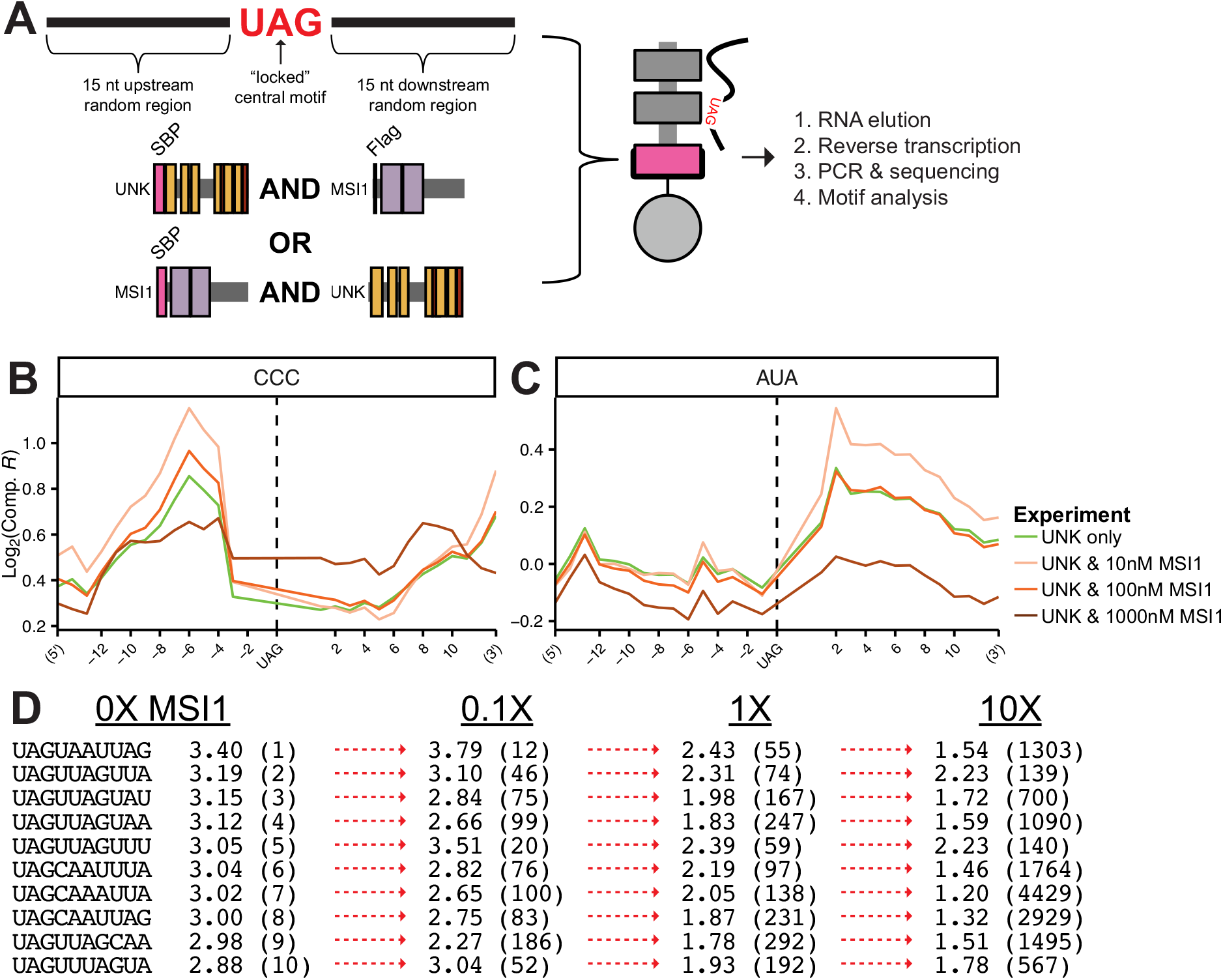
Competition of secondary motifs between MSI1 and UNK as determined via posCompRBNS. A) Design of posRNA and layout of posCompRBNS. B-C) Log_2_ posComp enrichments of B) CCC and C) AUA with UNK and MSI1 in competition at 0 (green), 10 (light orange), 100 (orange), or 1000 nM (dark orange). D) Rank depletion of the top ten UAGN7 10mers for UNK with MSI1 in competition. Arrows denote changes to rank with red denoting loss of top ten rank (new rank shown in parentheses next to enrichments).

Given the depth of posRBNS (discussed above) we calculated enrichment values for all 10mers starting with UAG (UAGN_7_). We compared both 10mer enrichments and ranks (e.g. 1 being the most enriched and so forth) with varying competitor concentrations. Indeed, all of UNK’s top 10mers (which are almost all bipartite UAGs) readily change in rank even at 0.1X MSI1 and drops in enrichments are seen at higher protein concentrations (**Figure 6D**). However, UNK’s top preferred 10mer (UAGUAAUUAG) is not as readily competed off (rank of 1 to 12) as MSI1 preferred motifs (UAGN_1-2_UAG). We noted that at the low MSI1 concentration 0.1X, UNK enrichments actually increase, we believe this effect to be related to low levels of competitors titrating away suboptimal UNK sequences thereby enhancing the enrichment of other sequences (similar to improving signal-to-noise).

When performing this assay in the opposite direction with SBP-tagged MSI1 and untagged UNK as competitor, we see that competition does not readily happen (**Supp. Figure 6C-F**). This is likely driven by MSI1’s greater affinity for RNA than that of UNK’s (**Figure 4A**). Very minor changes are observed for UNK’s secondary motifs (CCC and AUA) (**Supp. Figure 6C, E**) whereas UNK is unable to compete with MSI1 for bipartite UAG motifs (**Supp. Figure 6D**). When comparing 10mer rank across competition concentrations, very few large rank changes are observable where only three of the top ten drop out of the top ten at 0.1X and four drop at 1 to 10X UNK (**Supp. Figure 6F**). When taken together, these data highlight how sequence selectivity and affinity play into competitive binding aspects likely to carry over into *in vivo* binding. Previous studies have found that MSI1 is able to bind other RBP sequences better than those RBPs^30^ raising a perplexing question of how MSI1 achieves specificity *in vivo* (discussed below).

### MSI1 and UNK Demonstrate Some Overlapping Specificity *in vivo*

Understanding and modeling the biochemical details of how proteins bind RNA will allow for a deeper and much needed understanding of how targets are selected *in vivo*. However, to achieve this, longer natural sequences with evidence of *in vivo* binding need to be assessed *in vitro*. As noted above, we previously investigated species-specific UNK-RNA binding patterns between human and mouse using a massive scale *in vitro* assay involving thousands of natural RNA targets derived from iCLIP^33^. This approach enabled us to correlate our *in vitro* data with *in vivo* binding, binding strength, and regulation. We sought to apply this approach to examine how RBPs that target similar *in vitro* sequences (i.e. MSI1 and UNK) select for their targets *in vivo*. In this manner, we obtained iCLIP data for MSI1^37^ and UNK^11^. We defined binding sites in genes that were well-expressed in all the cell types in which iCLIP was performed (Methods, >5 transcripts per million (TPM)) and identified TAG-containing bound regions for MSI1 or UNK. We then compared these regions across proteins to identify conserved or protein-specific binding sites (**Supp. Figure 7A**).

At the transcript level, we identified 3,502 and 4,074 transcripts that were bound by MSI1 or UNK, respectively (Methods). Comparing these sites across proteins, we found that 50% of these transcripts are bound by both proteins (**Figure 7A**). Given the power of iCLIP at determining nucleotide-level resolution of binding, we were further able to determine where on each of the conserved transcripts MSI1 and UNK were bound. We largely found that due to each individual RBP’s regional binding preferences where MSI1 binds in the 3’UTR and UNK in the CDS preferentially^11,37^, 88% of the time, binding was differential where MSI1 bound to a UAG motif in the 3’UTR on the transcript whereas UNK binding was detected to a separate UAG motif in the CDS (**Figure 7B**). Interestingly, these data also show that neither protein binds UAG stop codons^11,37^, although why this is the case remains unclear.

**Figure 7.**
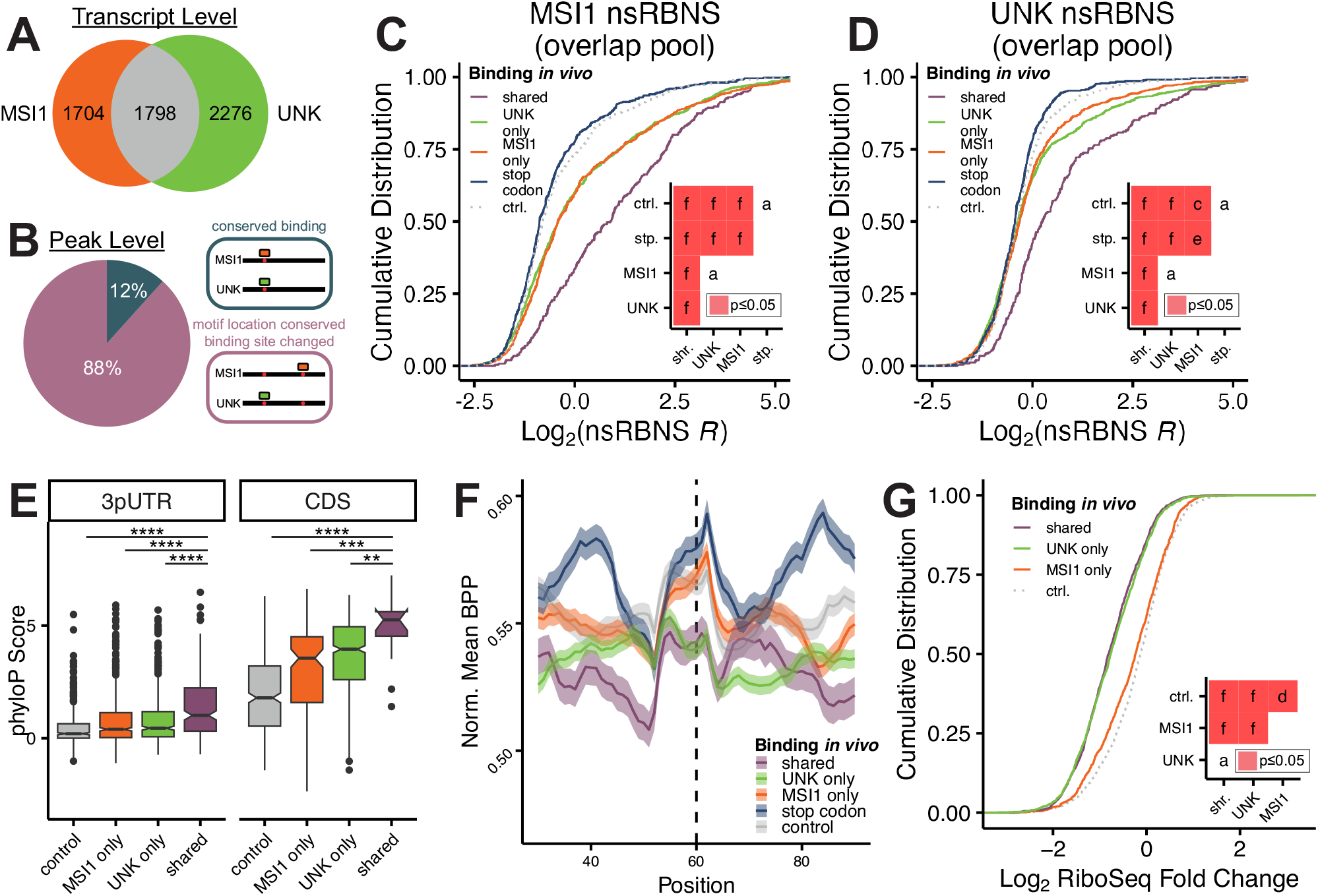
Analysis of MSI1 and UNK competition potential *in vivo* and *in vitro*. A) Transcript level conservation of iCLIP hits between MSI1 (from glioblastoma)^37^ and UNK (from SH-SY5Y or UNK-overexpressing HeLa)^11^. B) Motif level conservation of iCLIP hits between MSI1 (from glioblastoma)^37^ and UNK (from SH-SY5Y or UNK-overexpressing HeLa)^11^. C-D) Cumulative distribution function of log_2_ nsRBNS enrichments with C) MSI1 or D) UNK of all iCLIP hits: control (light grey; dotted), stop codon (dark blue), MSI1 only (orange), UNK only (green), and shared (purple) oligos. Insets show significance values for all comparisons via two-sided KS test and corrected for multiple comparisons via the BH procedure. Red denotes significant (p<=0.05). Values are as follows: a (ns), c (p<=0.05), e (p<=0.001), f (p<=0.0001). E) Box and whisker plot of the phyloP score of nsRBNS oligos: control (light grey), MSI1 only (orange), UNK only (green), and shared (purple). Center line denotes median (50th percentile) with bounds of box representing 25th to 75th percentiles and the whiskers denoting 5th to 95th percentiles. Outliers are denoted as individual points. Asterisks denote significance of each group compared to “shared” as determined via two-sided Wilcoxon test and are as follows: ** (p<=0.01), *** (p<=0.001), **** (p<=0.0001). F) Log_2_ fold change of mean base pair probability with a 10-nucleotide sliding window of the central region of shared, UNK only, MSI1 only, stop codon, and control oligos. Error bars show standard error of the mean. G) Cumulative distribution function of UNK RiboSeq^51^ fold change, log_2_, separated via iCLIP detection: control (light grey; dotted), MSI1 only (orange), UNK only (green), and shared (purple). Insets show significance values for all comparisons via two-sided KS test and corrected for multiple comparisons via the BH procedure. Red denotes significant (p<=0.05). Values are as follows: a (ns), d (p<=0.01), f (p<=0.0001).

To investigate protein-specific binding preferences and whether they are sequence or cellular context driven, we designed a natural sequence pool with all the overlapping regions as well as MSI1- and UNK-specific regions derived from iCLIP. We also included a large set of control sequences matched for UAG content and transcript region. We were further intrigued by the lack of UAG stop codon binding for both proteins^11,37^. To test if this is due to intrinsic features of stop codons or some other cellular factor, we included regions containing the stop codon for transcripts that displayed evidence of binding elsewhere. Finally, as a negative control we included mutant oligos where all occurrences of UAG were mutated to CCG to nullify binding. This resulted in a pool of 5,922 120nt naturally-derived sequences which we used to perform nsRBNS with MSI1 and UNK, separately. These assays were performed in duplicate with independent protein batches, and the replicates correlate well (**Supp. Figure 7B-C**).

As both MSI1 and UNK preferentially bind a UAG motif, we wanted to test the effects of mutating this motif to a CCG. As previously observed for UNK^11,33^, these mutations greatly diminish binding for both MSI1 and UNK (**Supp. Figure 7D-E**). Further, as the number of UAG motifs increase per oligo, so do the binding enrichments of both proteins (**Supp. Figure 7F-G**). These data confirm previously reported binding studies for MSI1 and UNK^27,35^ and demonstrate that this approach captures direct binding driven by known motifs even when embedded in the context of longer sequences.

Next, based on available iCLIP data^11,37^, we defined five classes of oligos: *control*, where no evidence of binding was observable for either protein; *stop codon*, stop codon regions from transcripts that MSI1 or UNK bound elsewhere on the transcript, *MSI1 only* or *UNK only*, where binding was observed for one protein but not both; and shared, where binding was observed for both proteins at the same region on the transcript. For both proteins, we observe that both control and stop codon oligos are lowly enriched (**Figure 7C-D**). Because both of these sets were designed to contain UAGs, this demonstrates that UAG alone is not sufficient to convey binding for either MSI1 or UNK. These data further demonstrate that the inability of MSI1 or UNK to bind the stop codon is in part due to an RNA sequence contribution rather than exclusively due to cellular context. Surprisingly, both MSI1 and UNK demonstrated fairly equivalent enrichments for MSI1 only or UNK only oligos (**Figure 7C-D**). These data suggest that while both proteins have distinct secondary motifs (**Figure 3B, D**) that allow for individual target selection and competition *in vitro* (**Figure 6**), *in vivo* binding patterns are not strictly defined by these sequences and rather are dependent on other cellular factors.

When looking at shared peaks across proteins, we observed the highest enrichments for both MSI1 and UNK (**Figure 7C-D**). These patterns are reminiscent of what we observed previously with species-specific binding patterns where highly conserved sequences were among the top bound^33^. To test that these may be highly conserved, we measured conservation (phyloP scores^64^) of the regions included in our pool and observed that shared oligos were the highest conserved (**Figure 7E**). We hypothesized that these higher enrichments may be due to increased accessibility to the binding motif; therefore, we performed *in silico* folding^61^, and compared the BPP of shared, UNK only, MSI1 only, stop codon, and control oligos. We found that shared oligos had the lowest BPP of the cohort especially immediately up and downstream of UAGs, likely enabling better tandem RBD engagement to unstructured RNA (**Figure 7F**). Further, we observed that the selected stop codon sequences had the highest BPP upstream and downstream, likely contributing to the lack of binding observed at these regions both *in vitro* and *in vivo*^11,37^. These data emphasize that overall motif accessibility plays an important role in RNA binding. This has been previously observed where many RBPs preferentially bind motifs present in hairpin loops rather than stems^27,29,62,65^.

To understand how these observed binding patterns reflect *in vivo* regulatory patterns, we utilized available RiboSeq data for MSI1 overexpressed in mouse cortical neuronal stem cells^52^ or UNK overexpressed in HeLa cells^51^ as both are translational regulators. We determined the translational impact of genes with shared or protein-specific peaks. For UNK, we see significant repression for shared and UNK only transcripts much greater than for MSI1 only or control transcripts (**Figure 7G**). For MSI1, we were unable to identify patterns that relate to binding, but this may be due to the fact that MSI1 may increase or decrease translation^66^, complicating this analysis (**Supp. Figure 7H**). Further, our data as well as previous studies have highlighted MSI1’s promiscuity when it comes to binding where MSI1 can often bind other RBP’s target RNAs better than those RBPs^30^. These findings suggest that it is not necessarily primary motif accessibility that dictates binding but rather a combination of regional sequences and protein stabilization that allow for RBP-RNA interactions.

## Discussion

RNA-protein interactions are critical for gene regulation. Disruption of these interactions can impact disease, as greater than 50% of genetic diseases can be linked to mutations within RNA regulatory elements that often interact with RNA-binding factors^67–69^. Despite this importance, RNA binding remains incompletely understood. Efforts have been made to catalog the RNA-binding preferences for the suite of RBPs^26–28,70^; however, much work remains to be done to cover all 1,500 RBPs within the human proteome^5^.

Here, we examine the RNA-binding preferences and overlapping specificities of three UAG-binding RBPs: MSI1, MSI2, and UNK. We find that these three proteins have significant motif overlap and while we identify unique specificities (**Figures 1-3**) our complete understanding of how these RNA-binding proteins select targets remains limited. Previous work by us and others have demonstrated that primary motifs alone are insufficient to define binding patterns with secondary motifs, RNA structure, and complex binidng being important drivers of interactions^33,71–74^. Our goal was to assess specificity as deeply as possible and to examine binding preferences and affinities domain-by-domain to understand how each domain alone and in tandem define interactions. When examining individual domains from MSI1, MSI2, and UNK, we found subtle differences that arise in sequence preference (**Figures 2 & 3**), perhaps allowing for RBP sequence selectivity (**Figure 6**). Importantly, more than half of RBPs contain either the same RBD type in tandem or a combination of multiple RBD types^5^ with striking examples like vigilin and ANKRD17 having more than ten domains^75–77^. While it is generally accepted that these multi-domain proteins have potential to bind and regulate RNA more tightly than their single domain counterparts^6,7,9–16^how these domains work together to impart target selection remains a challenging problem to solve.

Indeed, with MSI1, MSI2, and UNK we found that domains acting together bind RNA far better than any single domain alone, behaving as a “tandem anchoring” system (**Figures 4 & 5**). Understanding how intra-protein RBDs cooperate or antagonize each other in binding remains to be fully understood but will be important for predicting RBP targets and their related affinities. In the examples presented here, RBDs work together to enhance affinity, and as is shown for UNK, natural *in vivo* targets are poorly bound by individual domains (**Figure 5**). Multi-domain mechanisms in RNA-binding also require consideration of multiple binding sites on the RNA. It has been found that multiple binding sites in tandem can either aid in targeting of a given protein to the site, enhance specificity, avidity, and affinity of the interaction, or even result in competition between RBPs^27,32,78^. In the work presented here, we found that the presence of more than one binding site on a given RNA can enhance binding four-to ten-fold depending on which RBP is being assayed (**Figures 4 & 5**). In our massive-scale assays, natural binding sites have increased binding strength proportional to the number of motifs present.

An important question is how *in vitro* findings translate to an in vivo environment. Previous work by Lang et al. has demonstrated with enhanced CLIP (eCLIP) data that RBPs often form complexes to bind and regulate their RNA targets^32^.

We observe that MSI1 and UNK binding defined by iCLIP reveals approximately 50% transcript-level binding conservation, but far less binding-site conservation^11,37^. Even more, the specific transcript regions that are bound are different, with UNK preferring CDS and MSI1 preferring 3’ UTRs. Despite these clear differences observed *in vivo*, no significant differences are observed *in vitro* with nsRBNS. That is, *in vitro* UNK and MSI bound each other’s *in vivo* sites equally, highlighting that other factors outside of RNA sequence and structure largely contribute to protein-specific RNA binding (**Figure 7**). UNK is known to associate to ribosomes^51^ and this association may contribute to higher CDS binding *in vivo* than would be predicted based on *in vitro* data^33^.

Similar cellular context features have been observed previously to affect *in vivo* binding patterns. For instance, the LSm complex, which is essential for RNA degradation^79^, is recruited to histone mRNA by SLBP, where it’s association with histone mRNA is facilitated and enhanced through proteinprotein interactions between LSm4 and SLBP^80^. The RNA binding of IRP1 depends on intracellular iron levels where under low iron conditions, structured iron regulatory elements no longer compete with iron-sulfur clusters for binding to IRP1^81^. Expression patterns and localization of both the RBP and RNA also greatly impact binding^78^.

Another question is at how highly similar proteins (i.e. paralogs) often have independent functions. For instance, striking examples exist of paralogous proteins being highly conserved yet having differences in their binding preferences and regulation. For the paralogs hnRNPL and hnRNPLL, both proteins preferentially bind CA-rich motifs; however, hnRNPLL stringently prefers a spacing of two nucleotides whereas hnRNPL is more lenient^59^. Smith et al. further demonstrated that these differing specificities play a large part in how hnRNPL and hnRNPLL respond to RNA motif mutations^59^. For highly homologous (>70% conservation within the RBDs) such as the RBPs IGF2BP1 and IGF2BP2, slight differences within the KH domain variable loop result in 10-fold decreased binding of IGF2BP2 to IGF2BP1’s canonical sequence^82^. In our own work, we observe subtle but important binding differences between MSI1 and MSI2. While the two proteins are highly homologous (as discussed above), MSI2 possesses disordered domains at both termini, which can likely influence RNA binding or protein dimerization. Indeed, many previous studies have shown that RBP disordered regions can mediate RNA binding or dimerization^83–87^ (reviewed by McRae et al.^88^ and Kharel et al.^89^). Previous studies have shown that RRM1 of MSI2 dimerizes upon crystallization and that MSI1 can homodimerize or heterodimerize with MSI2 *in silico* and *in vivo*^90,91^. Therefore, while paralogous proteins are often highly similar, they are not identical and have been shown to have different RNA specificities, expression patterns, and protein-protein interactions.

The work described here presents a detailed, unbiased approach to fully catalog the RNA-binding preferences of multi-domain RBPs to better understand how individual RBPs specify for their target RNAs *in vivo*. Future work will involve building models starting with in vitro unbiased data wherein individual domains are catalogued independently and specificity is well-understood. However, adding *in vivo* features such as protein-protein interactions, RBP localization, and transcript region preferences will be essential to understand binding and regulation in cells.

## Methods

### Cloning of pGEX Constructs

pGEX-GST-SBP-MSI1 (mouse)^27^ and pGEX-GST-SBP-UNK (30-357)^27^ were gifted from Chris Burge (MIT). pGEX-GST-SBP-UNK ZnF1-3 and pGEX-GST-SBP-UNK ZnF4-6 have been previously published by our group^33^. DNA for MSI2 was ordered from Twist Biosciences. RBD sequences were selected for MSI1 as published by Iwaoka et al.^36^ and MSI2 as published by Lan et al.^90^. Domain annotations (**Supp. Table 1**) were verified with UniProt^54^ and InterPro^57^. Primers for insert amplification and plasmid construction were synthesized by Integrated DNA Technologies (IDT; **Supp. Table 2**).

Standard PCR was used to amplify plasmid inserts, restriction enzymes (New England Biolabs, NEB) were used to cleave plasmid backbones, and In-Fusion (Takara Bio) cloning was utilized according to manufacturer recommendations. Following In-Fusion cloning, plasmids were transformed into Stellar competent cells (Takara Bio) then miniprepped (Qiagen) and sequence verified via Sanger sequencing (Genewiz). Relevant plasmid information for this study (backbone, restriction sites used, insert, etc.) can be found in **Supp. Table 3**.

### Expression and Purification of Recombinant Proteins

SBP-MSI1 (mouse)^27^, SBP-MSI2, SBP-UNK (30-357)^27^, Flag-MSI1 (mouse), HIS-GST-UNK (30-357), SBP-MSI1 RRM1, SBP-MSI1 RRM2, SBP-MSI2 RRM1, SBP-MSI2 RRM2 SBP-UNK ZnF1-3^33^, and SBP-UNK ZnF4-6^33^, were purified as previously described^33^ with slight modifications. Rosetta E. coli competent cells (Novagen) were used for all purifications. Transformed cultures were grown in LB media at 37°C until an OD of 0.8 was reached, then induced with 0.5 mM IPTG (Thermo Scientific) at 16°C for 24 hours. Cultures were centrifuged at 4°C for 15 minutes at 4,000 x g to harvest cells, resuspended in lysis buffer (200 mM NaCl, 5 mM DTT, 50 mM HEPES, 3 mM MgCl_2_, 2 mM PMSF, 1 Pierce^TM^ protease inhibitor mini tablet/2 L; Thermo Scientific), sonicated, then incubated at 25°C for 30 minutes with 500 units/1 L culture Benzonase Nuclease (Sigma-Aldrich) and 5 units/1 L RQ1 RNase-free DNase (Promega). Lysate was clarified via centrifugation at 17,800 x g for 30 minutes at 4°C.

Pierce Glutathione Agarose (Thermo Scientific) was used for protein purification. Beads were equilibrated in low salt buffer (300 mM NaCl, 50 mM HEPES) prior to 1 hour incubation with lysate at 4°C. Recombinant protein-bound beads were washed in low salt buffer and high salt buffer (1 M NaCl, 50 mM HEPES) prior to incubation overnight at 4°C with 1:50 PreScission Protease (Cytiva) in cleavage buffer (20 mM HEPES, 100 mM NaCl, 5 mM DTT, 10% glycerol, 0.01% triton X-100). For SBP-MSI2, buffers were as follows due to predicted disordered regions: lysis buffer (200 mM NaCl, 5 mM DTT, 50 mM HEPES, 3 mM MgCl_2_, 2 mM PMSF, 1 Pierce^TM^ protease inhibitor mini tablet/2 L, 1% Triton X-100), low salt buffer (300 mM NaCl, 50 mM HEPES, 5 mM EDTA, 0.1% Triton X-100), and high salt buffer (1 M NaCl, 50 mM HEPES, 5 mM EDTA, 0.1% Triton X-100).

Proteins were eluted off the beads in excess cleavage buffer, concentrated via spin column (Vivaspin; Cytiva) and quality controlled via Pierce 660 nm assay (Thermo Scientific) and SDS-PAGE (4-12% gradient). For SBP-MSI1 (mouse), SBP-MSI2, SBP-UNK (30-357), Flag-MSI1 (mouse), HIS-GST-UNK (30-357), size exclusion chromatography (Superdex 200 Increase 10/300 GL; Cytiva) was used for further purification. Fractions were separated and collected in size exclusion buffer (20 mM HEPES, 1 M NaCl, 10 mM DTT, 0.01% triton X-100), and relevant fractions were pooled.

### Design and *in vitro* Transcription of RBNS Pools

20merRBNS and posRBNS DNA pool was synthesized by IDT (**Supp. Table 4**) and was *in vitro* transcribed using T7 RiboMAX Express large scale RNA kit (Promega) according to manufacturer protocols.

### Random RNA Bind-n-Seq (RBNS)

RBNS was performed as previously described^27,55^ with slight modifications. Recombinant SBP-tagged protein was incubated at 4°C for 30 minutes with MyOne Streptavidin T1 Dynabeads (Thermo Scientific) at multiple concentrations (SBP-MSI2: 50, 250 nM; SBP-MSI1 RRMs: 250, 500, 1000 nM; SBP-MSI2 RRMs: 250, 500, 1000 nM; SBP-UNK ZnF4-6: 250, 500, 1000 nM) in RBNS binding buffer (25 mM Tris, pH 7.5, 150 mM KCl, 3 mM MgCl_2_, 0.01% triton-X 100, 500 μg/mL BSA, 20 units/mL SUPERase·In (Thermo Fisher)). Unbound protein was removed, protein-bead complexes were resuspended in RBNS binding buffer, then incubated with 1 μM 20merRBNS RNA at 4°C for 1 hour. Protein-RNA-bead complexes were washed in RBNS wash buffer (25 mM Tris pH 7.5, 150 mM KCl, 0.01% triton X-100, 20 units/mL SUPERase·In) prior to elution at 60°C in RBNS elution buffer (0.1% SDS, 0.3 mg/mL proteinase K Thermo Fisher)) for 30 minutes. Elution was repeated and elutions were pooled prior to phenol chloroform extraction.

Eluted RNA was reverse transcribed with Superscript IV (Invitrogen) with RBNS RT primer (**Supp. Table 5**) and PCR amplified via Phusion DNA polymerase (New England Biolabs) with RBNS index primers and RBNS reverse primer (**Supp. Table 5**). Sequencing was performed on an Illumina NextSeq 1000. 6mer enrichments were calculated as the normalized count (frequency) of each 6mer within the bound pool versus the frequency in the input pool as reported previously^27^. Full binding reactions were performed in duplicate with independent protein batches, and the replicate enrichments were averaged. Logos were generated with the top 15 6mers, aligning to the top enriched 6mer. Fastqs for SBP-MSI1 and SBP-UNK were obtained from ENCODE^27^ (MSI1: ENCSR329RIP, UNK: ENCSR497VCL) and processed as detailed above. Fastqs for SBP-UNK ZnF1-3 and one replicate for SBP-UNK ZnF4-6 were obtained from GEO^33^ (GEO; GSE262560) and processed as detailed above. Final logos were plotted in RStudio^92^ (version 4.3.1) with R package ‘rkatss^93^‘ (version 0.0.0.9002).

### Positional RBNS (posRBNS)

posRBNS was performed similarly to RBNS (see above) with slight modifications. SBP-MSI1, SBP-MSI2, and SBP-UNK were incubated at 1, 10, 100, and 1000 nM with 1 μM posRBNS RNA. *K*mer enrichments were calculated positionally upstream and downstream of the central UAG where reads including an additional UAG outside of the central or the counted position were excluded. Logos were generated separately upstream and downstream with the top five 4mers and manually aligned. Final logos were plotted in RStudio^92^ with R package seqLogo^94^ (version 1.66.0).

### PosRBNS and iCLIP Pattern Analysis

MSI1 iCLIP-seq data from glioblastoma was downloaded from Uren et al.^37^ (GSE68800), MSI2 iCLIP-seq in K562 cells over-expressing MSI2 was downloaded from Karmakar et al.^38^ (GSE93210), and UNK iCLIP-seq data in HeLa cells overexpressing UNK or SH-SY5Y cells was downloaded from Murn et al.^11^ (E-MTAB-2279). As needed, peaks were converted from hg18 (MSI1) or hg19 (MSI2 and UNK) to hg38 using liftOver^95^ (version 1.24.0) in RStudio^92^ with liftOver chains obtained from UCSC. As all three proteins are cytoplasmically localized^11,44,46^, only peaks overlapping with exons were including. This was done with AnnotationHub^96^. Further, only one-to-one human-to-mouse orthologous genes with >5 TPM expression in glioblastoma, MSI2-overexpressing K562, SH-SY5Y, or UNK-overexpressing HeLa cells from the associated studies were included.

Peaks were expanded to 100 base pairs and sequences were obtained with function ‘getSeq’ using BSgenome.Hsapiens. UCSC.hg38^97^ and filtered for ‘TAG’ as both proteins primarily require a TAG to bind^11,27,35,37^. Peaks were centered around a ‘TAG’ motif and collapsed to 33 nucleotides. For MSI1 and MSI2, replicates were concatenated, and for UNK, both cell types were concatenated prior to downstream processing. Single protein peaks were sorted and merged with BEDTools^98^ (version 2.30). Peaks were expanded back out to 100 nucleotides to recenter at ‘TAG’ following the merge, then trimmed back down to 33 nucleotides with a central ‘TAG’ motif.

We annotated the genomic region of each peak according to the GENCODE (v34). Then we generated control unbound regions based on these genomic regions. For UTR regions, the control was randomly sampled from the same region of same gene. For CDS and intronic regions, the control was chosen among the regions that preserved the distance from the nearest splice sites.

Bipartite patterns were identified and counted in posRBNS and iCLIP reads and normalized for library size then to controls (input RNA for posRBNS and control regions for iCLIP).

### Fluorescence Polarization (FP)

3’ 6-FAM fluorescently-tagged RNA was ordered from IDT (Supp. Figure 6). Serial diluted recombinant SBP-tagged protein was incubated with 5 nM RNA in FP binding buffer (20 mM HEPES, 5 mM DTT, 137.5 mM NaCl, 0.01% triton X-100, 10 ng/μL BSA, 2 units/mL SUPERase·In) at 4°C for 15 minutes. For positional FP, 50 nM untagged random RNA (N_16_) was added to increase RNA competition. Following centrifugation at 1000 x g for 1 minute, FP was measured at 25°C with a CLARIOstar plate reader (BMG Labtech). FP binding assays were performed in triplicate with independent protein batches. Where relevant, delta FP was calculated to account for minimum and maximum FP. Data were fit in GraphPad Prism to a single site binding model to determine a K_d_.

### Natural Sequence RBNS (nsRBNS)

nsRBNS was performed similarly to RBNS (see above) with slight modifications. SBP-MSI1 and SBP-UNK were incubated at 100 nM with 1 μM nsRBNS RNA. SBP-UNK ZnF1-3 and SBP-UNK ZnF4-6 were assayed at 500 nM.

### nsRBNS Mapping and Enrichment Analysis

Reads were trimmed using fastx_toolkit^99^ (version 0.0.14) as needed prior to mapping with STAR^100^ (version 2.7.10b) with parameters set to –outFilterMultimapNMax 1 and –outFilterMismatchNmax 1. Seqk^101^ (version 2.3.0) was used to trim input fasta file. Enrichment was calculated as the frequency of the full-length sequence in the bound pool over the frequency in the input pool. For MSI1 and UNK nsRBNS, reads with less than 25 counts were excluded. Final sequences, enrichments, and relevant oligo information are included in **Supp. Data 1-4**.

### Competition Positional RBNS (compPosRBNS)

compPosRBNS was performed similarly to posRBNS (see above) with slight modifications. SBP-tagged proteins were held at 100 nM. Following removal of unbound SBP-tagged protein, untagged protein (UNK or Flag-MSI1) was added at 10, 100 or 1000 nM to incubate with protein-bead-RNA complexes for one hour. Elution, RT, and PCR was performed as described above. Enrichments were calculated positionally as detailed above then normalized to 0 nM antagonist protein.

### iCLIP and Peak Overlap Analysis

MSI1 and UNK iCLIP-seq data was processed as described above with slight modifications. Peaks were expanded out to 100 base pairs prior to concatenating replicates. Following merge, MSI1 and UNK peaks were intersected with BEDTools^98^ to determine overlapping regions. Multimapping peaks were removed. Peaks were expanded to 250 base pairs and aligned at a central ‘TAG.’ All overlapping peaks were included. For non-overlapping peaks, top-scoring iCLIP peaks were preferred with approximately 50:50 distribution between CDS and 3’UTR.

All CDS and 3’UTR sequences were downloaded from Ensembl BioMart^102^. Stop codon-centered sequences were selected for all overlapping peaks as well as top-scoring MSI1 or UNK bound genes. Controls were selected randomly from genes not demonstrating MSI1 or UNK binding via iCLIP and were matched for TAG content with bound regions. For iCLIP processing and pool assembly, R packages ‘AnnotationHub^96^,’ ‘dplyr^103^,’ ‘ensembldb^104^,’ ‘liftOver^95^,’ ‘seqParser^105^,’ ‘stringr^106^,’ and ‘tidyr^107^‘ were used.

## Acknowledgements

This work was in part supported by NIH T32 GM008570 (S.E.H.) and R35GM142864 (D.D.) as well as startup funds from UNC Chapel Hill to (D.D.).

## Author Contributions

Conceptualization, S.E.H. and D.D.; Methodology, S.E.H., K.B., B.B.G., J.G.M., and D.D.; Validation, S.E.H., K.B., B.B.G., J.G.M., and D.D.; Formal Analysis, S.E.H., Y.H., F.F.C., and D.D.; Investigation, S.E.H., Y.H., and D.D.; Resources, D.D.; Data Curation, S.E.H., Y.H., F.F.C., and D.D.; Writing – Original Draft, S.E.H. and D.D.; Writing – Review Editing, S.E.H., J.M., M.M.A., and D.D.; Visualization, S.E.H., Y.H., and D.D.; Supervision, M.A. and D.D.; Funding Acquisition, D.D.

