## Supplementary Information and Figures for "Dissecting RNA Selectivity Mediated by Tandem RNA-Binding Domains"

### Index of Supplementary Information for Harris *et al*.

Supplementary Figures


- Supp. Figure 1 → relates to Figure 1: Motif preferences for full-length MSI1, MSI2, and UNK as determined via RBNS
- Supp. Figure 2 → relates to Figure 2: Motif preferences for individual RBDs of MSI1, MSI2, and UNK as determined via RBNS
- Supp. Figure 3 → relates to Figure 3: Secondary motif, spacing, and base pair probability preferences for MSI1, MSI2, and UNK as determined via posRBNS
- Supp. Figure 4 → relates to Figure 4: Affinity of MSI1 and MSI2 for polyN RNA
- Supp. Figure 5 → relates to Figure 5: Combinatorial affinity of UNK for positional motifs
- Supp. Figure 6 → relates to Figure 6: Competition of secondary motifs between MSI1 and UNK as determined via posCompRBNS
- Supp. Figure 7 → relates to Figure 7: Analysis of MSI1 and UNK competition potential *in vivo* and *in vitro*


Supplementary Tables


- Supp. Table 1 → Domain regions for selected proteins
- Supp. Table 2 → Primer sequences for cloning of recombinant protein constructs
- Supp. Table 3 → Plasmid design for relevant plasmids
- Supp. Table 4 → Relevant oligo information for 20mer and posRBNS RNA
- Supp. Table 5 → Primer sequences for RBNS reverse transcription and PCR
- Supp. Table 6 → Sequences for synthetic fluorescence polarization RNA oligos


Supplementary References

Supplementary Download Items

- Supp. Data 1 → relates to Figure 5: UNK ZnF1-3 nsRBNS data table used for all analyses with sequences, relevant iCLIP information, enrichment values, relevant sequence information, and relevant oligo information
- Supp. Data 2 → relates to Figure 5: UNK ZnF4-6 nsRBNS data table used for all analyses with sequences, relevant iCLIP information, enrichment values, relevant sequence information, and relevant oligo information
- Supp. Data 3 → relates to Figure 7: MSI1 overlapping nsRBNS data table used for all analyses with sequences, relevant iCLIP information, enrichment values, relevant sequence information, and relevant oligo information
- Supp. Data 4 → relates to Figure 7: UNK overlapping nsRBNS data table used for all analyses with sequences, relevant iCLIP information, enrichment values, relevant sequence information, and relevant oligo information

#### SUPPLEMENTAL FIGURE LEGENDS

**Supp. Figure 1. Motif preferences for full-length MSI1, MSI2, and UNK as determined via RBNS.** A) Bar plot of $log_2$ RBNS enrichments of top 6mers for MSI1, MSI2, and UNK at all assayed concentrations. MSI1 and UNK data assayed by Dominguez *et al.*[1].

**Supp. Figure 2. Motif preferences for individual RBDs of MSI1, MSI2, and UNK as determined via RBNS.** A-C) Bar plot of $log_2$ RBNS enrichments of top 6mers for A) MSI1 RRM1 and MSI1 RRM2, B) MSI2 RRM1 and MSI2 RRM2, and C) UNK ZnF1-3 and UNK ZnF4-6 at all assayed concentrations. One replicate of UNK ZnF1-3 and UNK ZnF4-6 assayed by Harris *et al.*[2]. top ten 3mers and enrichment values as determined by iterative *k*mer analysis[1,3] included. D) Alignment of MSI1 RRM1 and MSI1 RRM2 as analyzed by BLAST[4]. RNA-contacting residues as determined by NMR structures[5,6] conserved across RRMs highlighted in red with similar amino acids highlighted in blue. E) Alignment of UNK ZnF1-3 and UNK ZnF4-6 as analyzed by BLAST[4]. RNA-contacting residues as determined by crystal structures[7] across domains highlighted in red with similar amino acids highlighted in blue and non-conserved residues highlighted in orange for ZnF1-3 and green for ZnF4-6.

**Supp. Figure 3. Secondary motif, spacing, and base pair probability preferences for MSI1, MSI2, and UNK as determined via posRBNS.** A) Heat map of the $log_2$ posRBNS enrichment of all 3mers for MSI1. B) Heat map of the base pair probability of all 3mers for MSI1. C) Heat map of the $log_2$ posRBNS enrichment of all 3mers for MSI2. D) Heat map of the base pair probability of all 3mers for MSI2. E) Heat map of the $log_2$ posRBNS enrichment of all 3mers for UNK. F) Heat map of the base pair probability of all 3mers for UNK. G-I) Line plot of the base pair probability of CCC for H) MSI1, I) MSI2, and J) UNK.

**Supp Figure 4. Affinity of MSI1 and MSI2 for polyN RNA.** A) Fluorescence polarization binding curves (n=3) for full-length MSI1 (orange circle), MSI1 RRM1 (light orange square), and MSI1 RRM2 (dark orange triangle) incubated with a polyN oligo. Data are presented as mean values +/- SD. B) Fluorescence polarization binding curves (n=3) for full-length MSI2 (teal circle), MSI2 RRM1 (light teal square), and MSI2 RRM2 (dark teal triangle) incubated with a polyN oligo. Data are presented as mean values +/- SD. C) Fluorescence polarization binding curves (n=3) for all RNA-binding domains of UNK (green circle), UNK ZnF1-3 (light green square), and UNK ZnF4-6 (dark green triangle) incubated with a polyN oligo. Data are presented as mean values +/- SD.

**Supp. Figure 5. Combinatorial affinity of UNK for positional motifs.** A) Fluorescence polarization binding curves (n=3) for UNK incubated with a polyU oligo (blue circle) or a polyC oligo (yellow square). Data are presented as mean values +/- SD. B) Correlation plot $\log_2$ ZnF1-3 nsRBNS enrichments versus $\log_2$ UNK nsRBNS enrichments. Pearson's correlation coefficient and p value included. C) Correlation plot $\log_2$ ZnF4-6 nsRBNS enrichments versus $\log_2$ UNK nsRBNS enrichments. Pearson's correlation coefficient and p value included. D) Correlation plot $\log_2$ ZnF1-3 nsRBNS enrichments versus $\log_2$ ZnF4-6 nsRBNS enrichments. Pearson's correlation coefficient and p value included.

**Supp. Figure 6. Competition of secondary motifs between MSI1 and UNK as determined via posCompRBNS.** A-B) $\text{Log}_2$ posComp enrichments of A) CCC or B) AUA with UNK and MSI1 in competition at 0, (green), 10 (light orange), 100 (orange), or 1000 nM (dark orange). Only oligos with a single UAG were considered. C-E) $\text{Log}_2$ posComp enrichments of C) CCC, D) UAG, E) AUA with MSI1 and UNK in competition at 0, (orange), 10 (light green), 100 (green), or 1000 nM (dark green). F) Rank depletion of the top ten $\text{UAGN}_7$ 10mers for MSI1 with UNK in competition. Arrows denote changes to rank with grey denoting maintenance within top ten and red denoting loss of top ten rank.

**Supp. Figure 7. Analysis of MSI1 and UNK competition potential *in vivo* and *in vitro*.** A) Diagram of iCLIP peak overlap analysis method between MSI1 (from glioblastoma)[8] and UNK (from SH-SY5Y or UNK-overexpressing HeLa)[9]. B-C) Correlation plot of two experimental B) MSI1 or C) UNK overlap pool nsRBNS replicates. Pearson's correlation coefficient and p value included. D-E) Scatter plot of $\log_2$ D) MSI1 or E) UNK overlap pool nsRBNS enrichments of wild-type (Y-axis) versus total motif mutant (X-axis) oligos. $\text{Log}_2$ change in enrichment (wt-mut) was calculated for each sequence pair: > 0.5 defined as bound better in wt (blue), < -0.5 defined as bound better in mut (red), 0 +/- 0.5 defined as similar binding (grey). Significance determined via paired, one-sided Wilcoxon test. F-G) Cumulative distribution function of $\log_2$ F) MSI1 or G) UNK overlap pool nsRBNS enrichment of all oligos separated by UAG motif content. Insets show significance values for all comparisons via two-sided KS test and corrected for multiple comparisons via the BH procedure. Red denotes significant ($p \leq 0.05$), and grey denotes nearing significant ($p \leq 0.1$). Values are as follows: a (ns), b ($p \leq 0.1$), d ($p \leq 0.01$), e ($p \leq 0.001$), f ($p \leq 0.0001$). H) Cumulative distribution function of MSI1 RiboSeq[10] absolute fold change, $\log_2$, separated via iCLIP detection: control (light grey; dotted), MSI1 only (orange), UNK only (green), and shared (purple). Insets show significance values for all comparisons via two-sided KS test and corrected for multiple comparisons via the BH procedure. Red denotes significant ($p \leq 0.05$). Values are as follows: a (ns), c ($p \leq 0.05$), d ($p \leq 0.01$), e ($p \leq 0.001$), f ($p \leq 0.0001$).

#### SUPPLEMENTARY TABLES

##### Supp. Table 1. Domain regions for selected proteins

| Protein | Domain | Start | Stop |
|---|---|---|---|
| MSI1 | RRM | 20 | 103 |
| MSI1 | RRM | 109 | 191 |
| MSI2 | Disordered | 1 | 28 |
| MSI2 | RRM | 21 | 111 |
| MSI2 | RRM | 110 | 187 |
| MSI2 | Disordered | 264 | 286 |
| UNK | Disordered | 1 | 24 |
| UNK | Disordered | 1 | 24 |
| UNK | ZnF | 34 | 73 |
| UNK | ZnF | 84 | 113 |
| UNK | ZnF | 124 | 154 |
| UNK | ZnF | 215 | 240 |
| UNK | Disordered | 239 | 265 |
| UNK | ZnF | 251 | 285 |
| UNK | ZnF | 293 | 321 |
| UNK | Disordered | 324 | 343 |
| UNK | Coil | 647 | 681 |
| UNK | Coil | 696 | 723 |

##### Supp. Table 2. Primer sequences for cloning of recombinant protein constructs

| Primer name | Sequence |
|---|---|
| pGEX-GST-Flag-MSI1 (mouse) forward primer / pGEX-GST-SBP-MSI1 swap (mouse) forward primer | GCGTGAACCGGGATCCATGGAGACTGACGCGC |
| pGEX-GST-Flag-MSI1 (mouse) reverse primer | ACGATGCGGCCGCCTATCAGTGGTACCCATTG |
| pGEX-GST-SBP-MSI1 swap (mouse) mid1 forward primer | CTCGCCGCACGACCCCAAGAAGATCTTCGTGG |
| pGEX-GST-SBP-MSI1 swap (mouse) mid1 reverse primer | CCACGAAGATCTTCTTGGGGTCGTGCGGCGAG |
| pGEX-GST-SBP-MSI1 swap (mouse) mid2 forward primer | AAAGGAGGTGATGTCCATGGTCACTCGGACGTG CAAGATGTTCATCG |
| pGEX-GST-SBP-MSI1 swap (mouse) mid2 reverse primer | CGATGAACATCTTGCACGTCCGAGTGACCATGG ACATCACCTCCTTT |
| pGEX-GST-SBP-MSI1 swap (mouse) mid3 forward primer | AAGAGCACAGCCTAAGCCGACAGGCTCAGCCC |
| pGEX-GST-SBP-MSI1 swap (mouse) mid3 reverse primer | GGGCTGAGCCTGTCGGCTTAGGCTGTGCTCTT |
| pGEX-GST-SBP-MSI1 swap (mouse) reverse primer | GTCACGATGCGGCCGCTCAGTGGTACCCATTG |

| | |
|---|---|
| pGEX-GST-SBP-MSI1 RRM1 (mouse) forward primer | GCGTGAACCGGGATCCTGCAAGATGTTCATCG |
| pGEX-GST-SBP-MSI1 RRM1 (mouse) reverse primer | ACGATGCGGCCGCTCACTTAGGCTGTGCTCTT |
| pGEX-GST-SBP-MSI1 RRM2 (mouse) forward primer | GCGTGAACCGGGATCCAAGAAGATCTTCGTGG |
| pGEX-GST-SBP-MSI1 RRM2 (mouse) reverse primer | ACGATGCGGCCGCTCAGGACATCACCTCCTTT |
| pGEX-GST-SBP-MSI2 forward primer | GCGTGAACCGGGATCCATGGAGGCAAATGGGA |
| pGEX-GST-SBP-MSI2 reverse primer | GTCACGATGCGGCCGCTCAATGGTATCCATTT |
| pGEX-GST-SBP-MSI2 RRM1 forward primer | GCGTGAACCGGGATCCGGTAAAATGTTTATCG |
| pGEX-GST-SBP-MSI2 RRM1 reverse primer | ACGATGCGGCCGCTCATTTCTTTGTTCTTGTG |
| pGEX-GST-SBP-MSI2 RRM2 forward primer | GCGTGAACCGGGATCCAAGAAAATATTTGTAG |
| pGEX-GST-SBP-MSI2 RRM2 reverse primer | ACGATGCGGCCGCTCACGGCTGAGCTTTCTTA |
| pGEX-GST-SBP-UNK swap forward primer / pGEX-GST-SBP-UNK dual forward primer | GCGTGAACCGGGATCCCCTCGGTGGCAAGAGA |
| pGEX-GST-SBP-UNK swap mid1 forward primer / pGEX-GST-SBP-UNK dual mid1 forward primer | TTCCTCAGCTGTGTCCGGCCAGACCACGGTAG |
| pGEX-GST-SBP-UNK swap mid1 reverse primer / pGEX-GST-SBP-UNK dual mid1 reverse primer | CTACCGTGGTCTGGCCGGACACAGCTGAGGAA |
| pGEX-GST-SBP-UNK swap mid2 forward primer | GATCCTCAGCGAGGAGCCGCAGCACTACACGT |
| pGEX-GST-SBP-UNK swap mid2 reverse primer | ACGTGTAGTGCTGCGGCTCCTCGCTGAGGATC |
| pGEX-GST-SBP-UNK swap mid3 forward primer | GGAGGCCTTGCAGAATAGCCCCACCCAGCCAG |
| pGEX-GST-SBP-UNK swap mid3 reverse primer | CTGGCTGGGTGGGGCTATTCTGCAAGGCCTCC |
| pGEX-GST-SBP-UNK swap reverse primer / pGEX-GST-SBP-UNK dual reverse primer | ACGATGCGGCCGCTCAGCTCACAGGCACCGAG |

**Supp. Table 3. Plasmid design for relevant plasmids**

| Backbone | Insert | Tag(s) | Restriction Sites |
|---|---|---|---|
| pGEX-6P-1 | MSI1 (mouse) | GST, Flag | XhoI, NotI |
| pGEX-6P-1 | MSI1 swap (mouse; 1-19,109-191,104-108,20-103,192-362) | GST, SBP | BamHI, NotI |
| pGEX-6P-1 | MSI1 RRM1 (mouse; 20-103) | GST, SBP | BamHI, NotI |
| pGEX-6P-1 | MSI1 RRM2 (mouse; 109-191) | GST, SBP | BamHI, NotI |
| pGEX-6P-1 | MSI2 | GST, SBP | BamHI, NotI |
| pGEX-6P-1 | MSI2 RRM1 (21-111) | GST, SBP | BamHI, NotI |

| | | | |
|---|---|---|---|
| pGEX-6P-1 | MSI2 RRM2 (110-187) | GST, SBP | BamHI, NotI |
| pGEX-6P-1 | UNK (30-357) | HIS, GST | XhoI, NotI |
| pGEX-6P-1 | UNK swap (204-335,175-203,31-174,336-357) | GST, SBP | BamHI, NotI |
| pGEX-6P-1 | UNK dual (204-335, 165-357) | GST, SBP | BamHI, NotI |

**Supp. Table 4. Relevant oligo information for 20mer and posRBNS RNA**

| Oligo Name | Sequence |
|---|---|
| 20merRBNS DNA template | CCTTGACACCCGAGAATTCCANNNNNNNNNNNNNNNNNNNN<br>GATCGTCGGACTGTAGAACTCCCTATAGTGAGTCGTATTA |
| posRBNS DNA template | CCTTGACACCCGAGAATTCCANNNNNNNNNNNNNNNCTANN<br>NNNNNNNNNNNNNGATCGTCGGACTGTAGAACTCCCTATAGT<br>GAGTCGTATTA |
| 20merRBNS RNA | GAGUUCUACAGUCCGACGAUCNNNNNNNNNNNNNNNNNNN<br>NUGGAAUUCUCGGGUGUCAAGG |
| posRBNS RNA | GAGUUCUACAGUCCGACGAUCNNNNNNNNNNNNNNNUAGN<br>NNNNNNNNNNNNNNUGGAAUUCUCGGGUGUCAAGG |

**Supp. Table 5. Primer sequences for RBNS reverse transcription and PCR**

| Primer name | Sequence |
|---|---|
| RBNS RT primer | TCCTTGGCACCCGAGAATTCCANNNNNNNNNTGATGCTCAATCCT<br>CTGTTG |
| RBNS index primer | CAAGCAGAAGACGGCATACGAGAT[$N_6$<br>barcode]GTGACTGGAGTTCCTTGGCACCCGAGAATTCCA |
| RBNS reverse primer | AATGATACGGCGACCACCGAGATCTACACGTTCAGAGTTCTACAG<br>TCCGA |

**Supp. Table 6. Sequences for synthetic fluorescence polarization RNA oligos**

| Oligo name | Sequence |
|---|---|
| Mono-UAG FP RNA | UAGUUUAGUUUAGUU /6-FAM/ |
| Dual-UAG FP RNA | UUAGUUAGUU /6-FAM/ |
| Tri-UAG FP RNA | AAAAAAAAUUAGAAAAAA /6-FAM/ |
| CCC-UAG FP RNA | CCCCCCAAUUAGAAAAAA /6-FAM/ |
| UAG-AUA FP RNA | AAAAAUUAGUUAAUA /6-FAM/ |
| CCC-UAG-AUA FP RNA | CCCCCCAAUUAGUUAAUA /6-FAM/ |
| polyN FP RNA | NNNNNNNNNNNNNNNN /6-FAM/ |
| polyC FP RNA | CCCCCCCCCCCCCCCC /6-FAM/ |
| polyU FP RNA | UUUUUUUUUUUUUUUU /6-FAM/ |

#### Supp. Figure 1

A

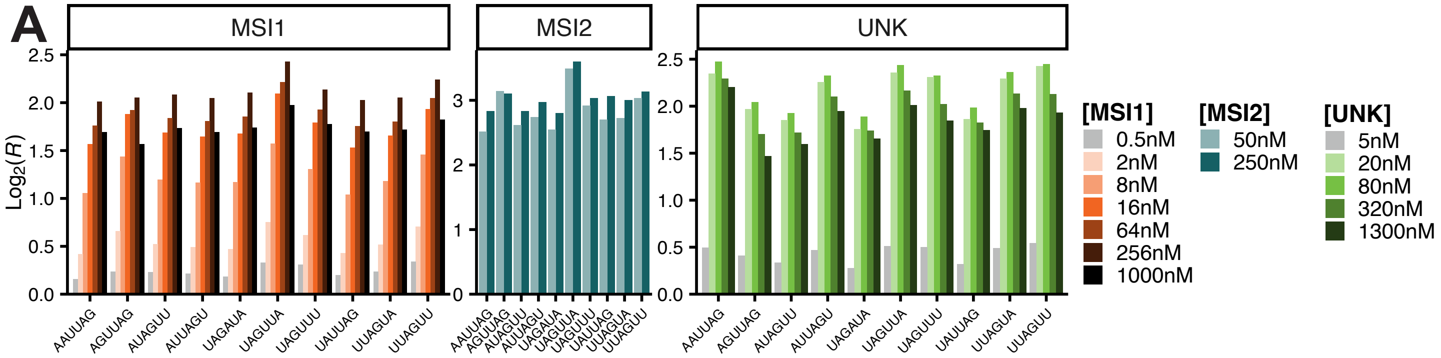

### Supp. Figure 2

**A**

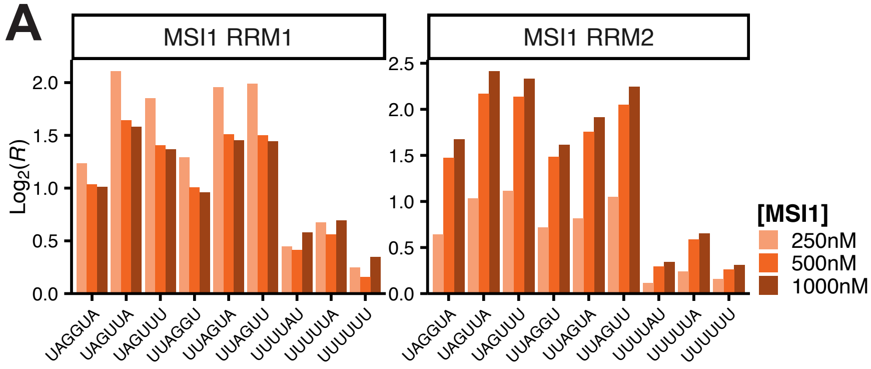


| RRM1 | | RRM2 | |
|---|---|---|---|
| UAG | 2.20 | UAG | 2.35 |
| UUA | 1.29 | UUA | 1.25 |
| UAU | 1.19 | UAA | 1.15 |
| AUA | 1.14 | UAU | 1.12 |
| CCC | 1.13 | AUA | 1.08 |
| UAA | 1.12 | CCC | 1.07 |
| CUA | 1.09 | GUA | 1.07 |
| AGU | 1.08 | AUU | 1.05 |
| UAC | 1.06 | UUG | 1.05 |
| GUA | 1.26 | GUU | 1.06 |

**B**

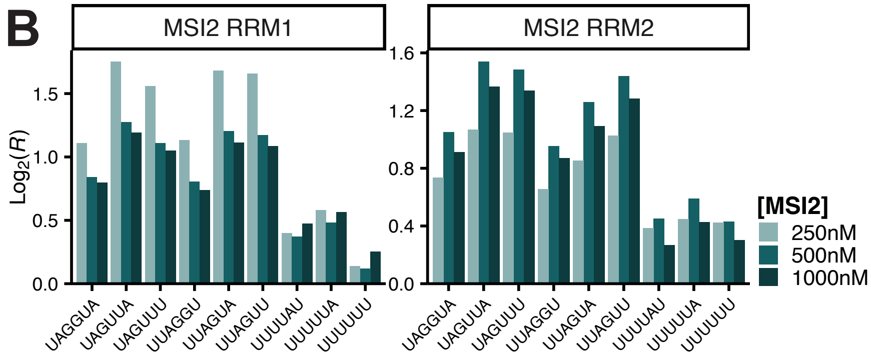


| RRM1 | | RRM2 | |
|---|---|---|---|
| UAG | 1.92 | UAG | 1.22 |
| UUA | 1.33 | UUA | 1.17 |
| AUA | 1.25 | UAU | 1.13 |
| UAA | 1.19 | AUU | 1.09 |
| AGU | 1.15 | GUA | 1.07 |
| UAU | 1.15 | UUC | 1.07 |
| CUA | 1.11 | GUU | 1.08 |
| CAG | 1.08 | CCC | 1.06 |
| UAC | 1.08 | UUG | 1.06 |
| GUA | 1.33 | CUU | 1.09 |

**C**

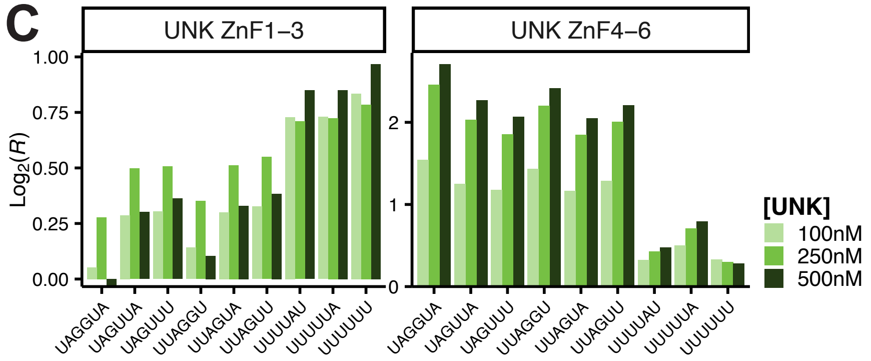


| ZnF1-3 | | ZnF4-6 | |
|---|---|---|---|
| UUA | 1.25 | UAG | 2.98 |
| UAU | 1.20 | UUA | 1.34 |
| CUU | 1.17 | UAU | 1.29 |
| AUU | 1.12 | AUA | 1.23 |
| UUC | 1.11 | UAA | 1.16 |
| CCC | 1.09 | CUA | 1.12 |
| UCU | 1.09 | AUU | 1.09 |
| UGU | 1.08 | AAA | 1.10 |
| CUA | 1.08 | UAC | 1.09 |
| AUA | 1.08 | GUA | 1.50 |

**D**

```
MSI1 RRM1: KMFIGGLSWQTTQEGLREYFGQFGEVKECLVMRDPLTKRSR
           K+F+GGLS   TT  E  +++YF  QFG+V  +  ++M  D   T  R  R
MSI1 RRM2: KIFVGGLSVNTTVEDVKQYFEQFGKVDDAMLMFDKTTNRHR

R1: GFGFVTFMDQAGVDKVLAQSRHELDSKTIDPKVAFPRRAQPK
    GFGFVTF   +   V+KV      HE+++K  ++  K       +AQPK
R2: GFGFVTFESEDIVEKVCEIHFHEINNKMVECK-----KAQPK
```

**E**

```
UNK ZnF1-3: YTCFHWHFVNQRRRRSIRRRDGTFNYSPDVYCTKYDEATGLCPEG
            Y  C  ++H    ++  RRRS  R+       +    P+V            G  C   G
UNK ZnF4-6: YACPYYH-NSKDRRRSPRKHKYRSSPCPNVKHGDEWGDPGKCENG

ZnF1-3: DECPFLHRTTGDTERRYHLRYYKTGICIHETDSKGNCTKNGLHCAFAH
        D  C  +  H       TE+++H    YK+   C  ++      G+C  +  G   CAFAH
ZnF4-6: DACQYCHTR---TEQQFHPEIYKSTKC-NDMQQSGSCPR-GPFCAFAH
```

### Supp. Figure 3

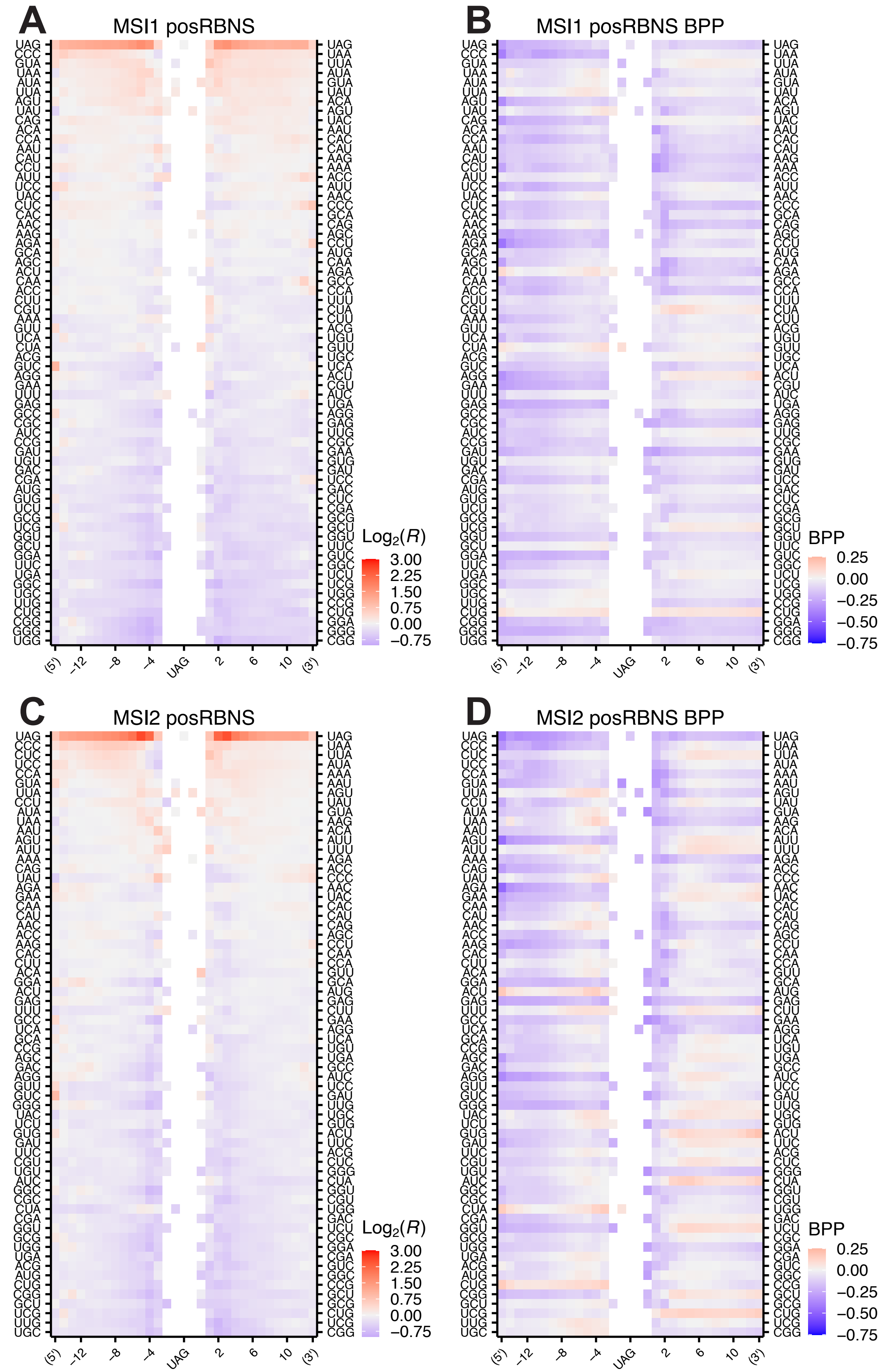

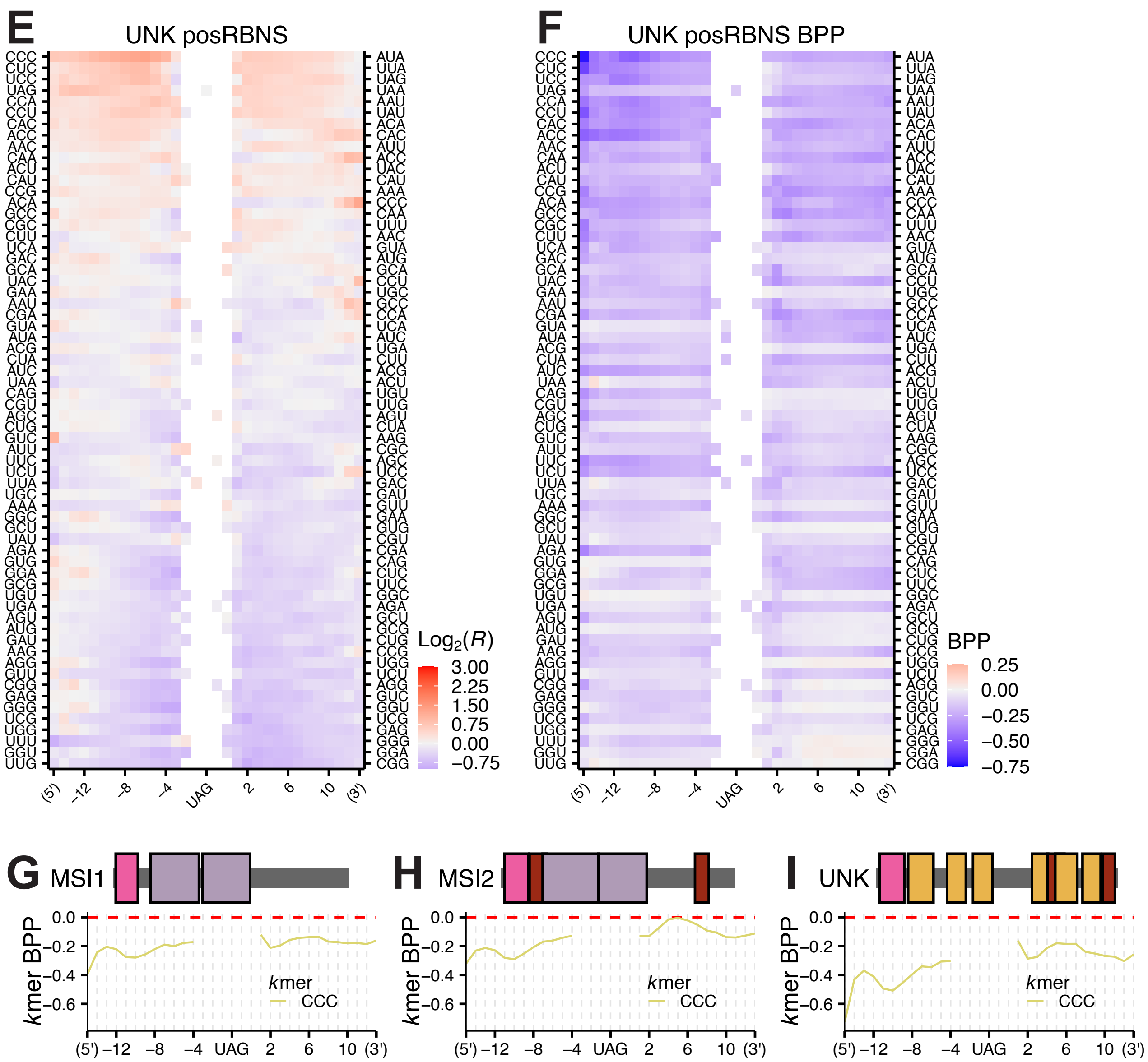
E
UNK posRBNS
CCC CUC UCC UAG CCA CCU CAC ACC AAC CAA ACU CAU CCG ACA GCC CGC CUU UCA GAC GCA UAC GAA AAU CGA GUA AUA ACG CUA AUC UAA CAG CGU AGC CUG GUC AUU UUC UCU UUA UGC AAA GGC GCU UAU AGA GUG GGA GCG UGU UGA AGU AUG GAU AAG AGG GUU CGG GAG GGG UCG UGG UUU GGU UUG
AUA UUA UAG UAA AAU UAU ACA CAC AUU ACC UAC CAU AAA CCC CAA UUU AAC GUA AUG GCA CCU UGC GCC CCA UCA AUC UGA CUU ACG ACU UGU UUG AGU CUA AAG CGC AGC UCC GAC GAU GUU GAA GUG CGU CGA CAG CUC UUC GGC AGA GCU GCG CUG CCG UGG UCU AGG GUC GGU UCG GAG GGG GGA CGG
(5') -12 -8 -4 UAG 2 6 10 (3')
Log2(R)
3.00 2.25 1.50 0.75 0.00 -0.75
F
UNK posRBNS BPP
CCC CUC UCC UAG CCA CCU CAC ACC AAC CAA ACU CAU CCG ACA GCC CGC CUU UCA GAC GCA UAC GAA AAU CGA GUA AUA ACG CUA AUC UAA CAG CGU AGC CUG GUC AUU UUC UCU UUA UGC AAA GGC GCU UAU AGA GUG GGA GCG UGU UGA AGU AUG GAU AAG AGG GUU CGG GAG GGG UCG UGG UUU GGU UUG
AUA UUA UAG UAA AAU UAU ACA CAC AUU ACC UAC CAU AAA CCC CAA UUU AAC GUA AUG GCA CCU UGC GCC CCA UCA AUC UGA CUU ACG ACU UGU UUG AGU CUA AAG CGC AGC UCC GAC GAU GUU GAA GUG CGU CGA CAG CUC UUC GGC AGA GCU GCG CUG CCG UGG UCU AGG GUC GGU UCG GAG GGG GGA CGG
(5') -12 -8 -4 UAG 2 6 10 (3')
BPP
0.25 0.00 -0.25 -0.50 -0.75
G MSI1
H MSI2
I UNK
kmer BPP
0.0 -0.2 -0.4 -0.6
kmer
CCC
(5') -12 -8 -4 UAG 2 6 10 (3')

#### Supp. Figure 4

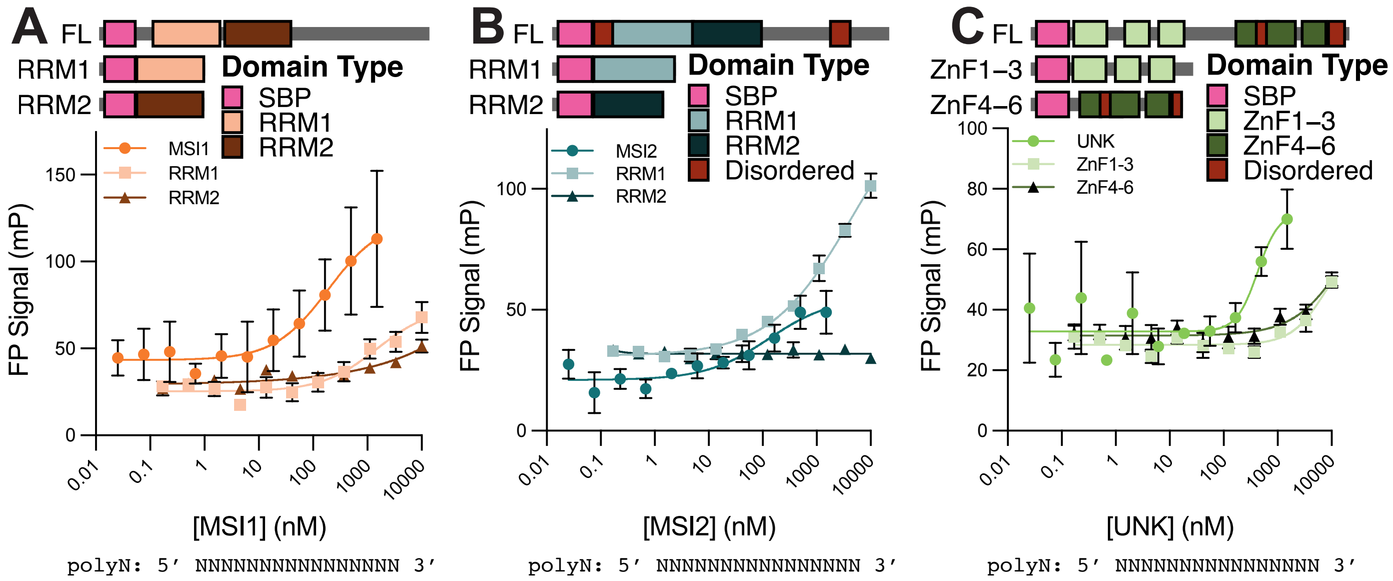

#### Supp. Figure 5

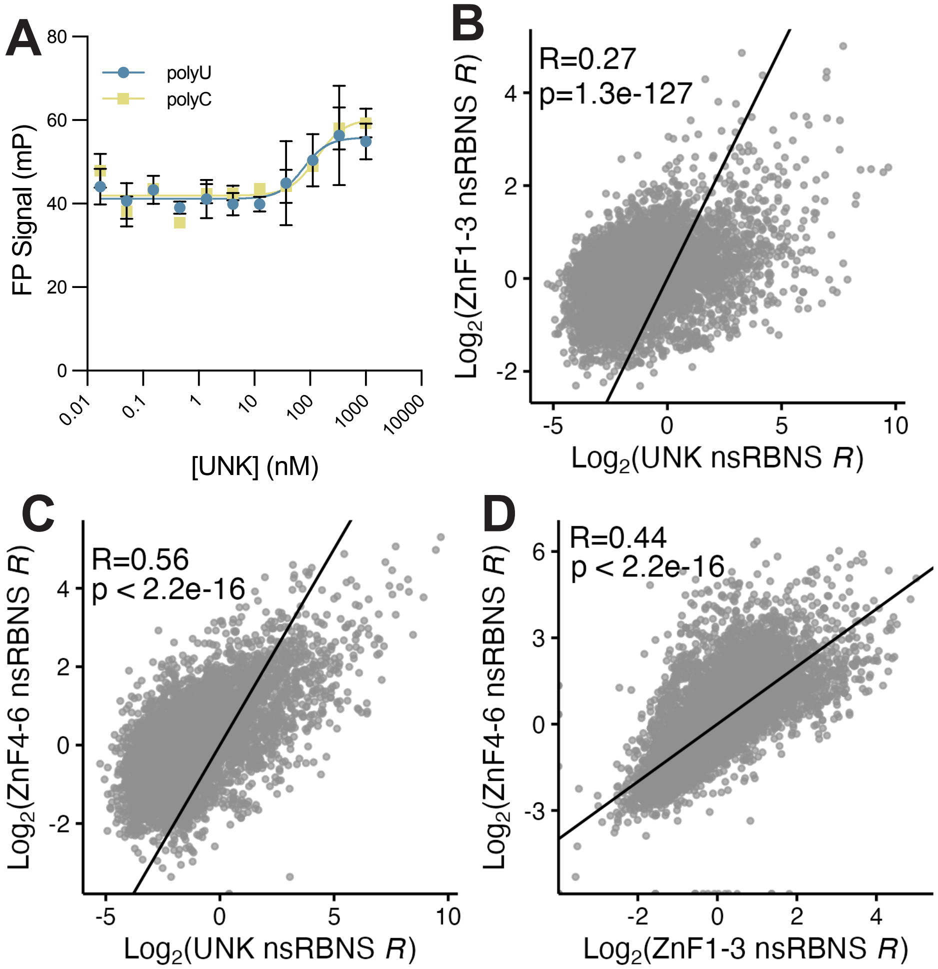

### Supp. Figure 6

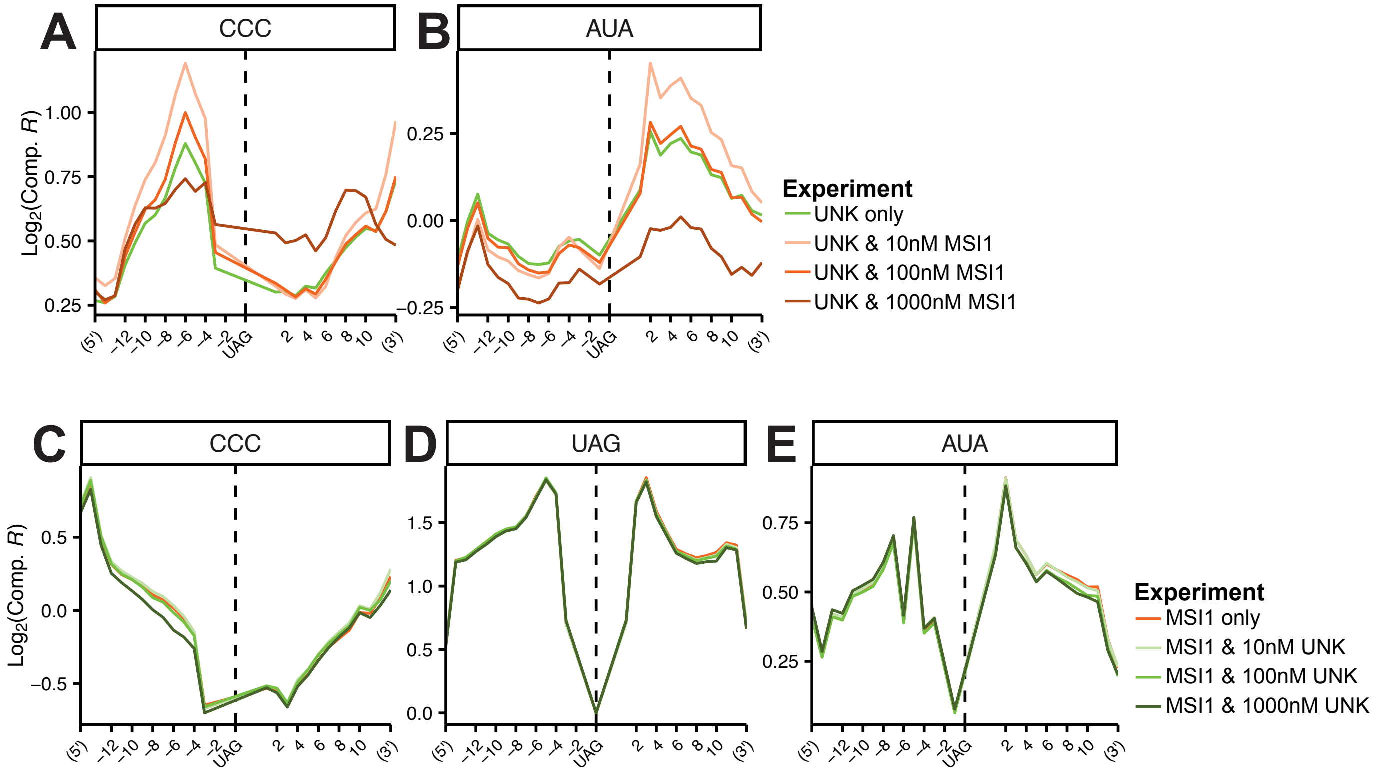


**F**

| | 0X UNK | 0.1X | 1X | 10X |
|---|---|---|---|---|
| UAGUUAGUUA | 4.77 (1) | 8.06 (1) | 8.11 (1) | 8.13 (1) |
| UAGUUAGUAG | 4.52 (2) | 7.02 (4) | 7.67 (3) | 7.44 (3) |
| UAGUUAGUGU | 4.44 (3) | 6.02 (17) | 6.26 (14) | 6.30 (13) |
| UAGUUAGUAU | 4.42 (4) | 6.98 (5) | 6.96 (6) | 6.88 (7) |
| UAGUUUAGCA | 4.41 (5) | 6.56 (9) | 6.41 (12) | 6.53 (10) |
| UAGUUUAGUA | 4.40 (6) | 7.47 (3) | 7.49 (4) | 7.42 (4) |
| UAGUUAGUUU | 4.29 (7) | 6.96 (6) | 6.89 (7) | 6.85 (9) |
| UAGUUAGCAU | 4.18 (8) | 5.97 (22) | 6.17 (16) | 6.03 (18) |
| UAGUUAGUAA | 4.18 (9) | 6.78 (8) | 6.69 (9) | 6.92 (6) |
| UAGUUAGCAA | 4.01 (10) | 5.81 (29) | 5.68 (31) | 5.63 (32) |

### Supp. Figure 7

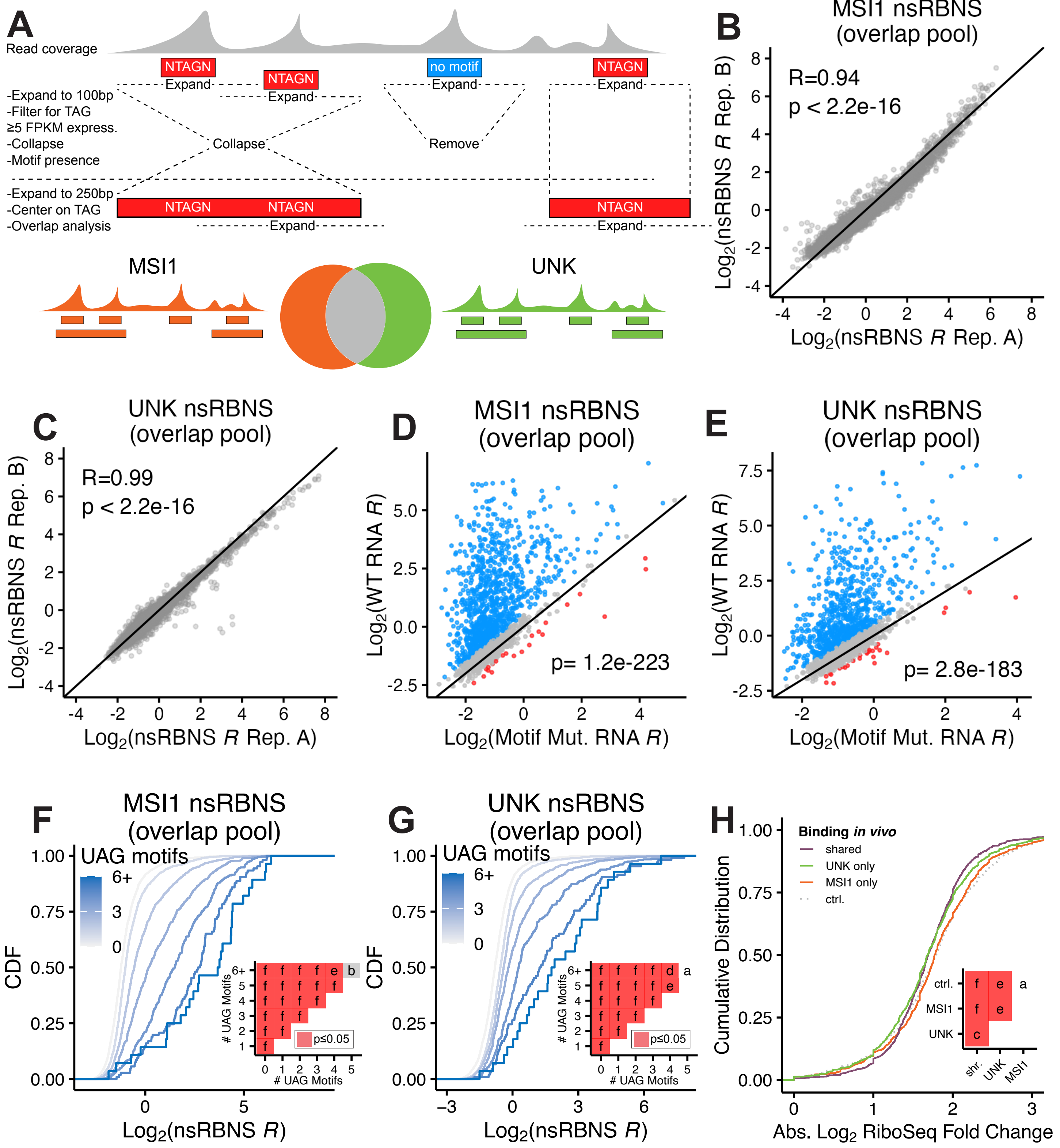
